# A member of a new dynamin superfamily modulates mitochondrial membrane branching in *Trypanosoma brucei*

**DOI:** 10.1101/2023.06.28.546890

**Authors:** Chloé Alexandra Morel, Corinne Asencio, David Moreira, Corinne Blancard, Bénédicte Salin, Etienne Gontier, Stéphane Duvezin-Caubet, Manuel Rojo, Frédéric Bringaud, Emmanuel Tetaud

## Abstract

Unlike most other eukaryotes, where mitochondria continuously fuse and divide, the mitochondrion of trypanosome cells forms a single and continuously interconnected network that divides only during cytokinesis. However, the machinery governing mitochondrial remodeling and interconnection of trypanosome mitochondrion remain largely unknown. We functionally characterize a novel dynamin-superfamily protein (DSP) from *T. brucei* (*Tb*MfnL) which shares close similarity with a family of homologs present in various eukaryotic and prokaryotic phyla, but not in opisthokonts like mammals and budding yeast. The sequence and domain organization of *Tb*MfnL is distinct and it is phylogenetically very distant from the yeast and mammalian dynamin-related proteins involved in mitochondrial fusion/fission dynamincs, such as Opa1 and Mfn. *Tb*MfnL localizes to the inner mitochondrial membrane facing the matrix, and upon overexpression, induces a strong increase in the interconnection and branching of mitochondrial filaments in a GTPase-dependent manner. *Tb*MfnL is a component of a novel membrane remodeling machinery with an unprecedented matrix-side localization that is able to modulate the degree of intermitochondrial connections.

## INTRODUCTION

*Trypanosoma brucei* is a parasite responsible for African sleeping sickness as well as the related cattle disease Nagana, affecting sub-Saharan Africa. *T. brucei* is transmitted to mammals by an insect vector: the tsetse fly (*Glossina* spp.). During the trypanosome life cycle, the parasite exists in at least two replicative forms; the procyclic form (PCF), transmitted by the tsetse fly, and the bloodstream form (BSF) responsible for diseases in vertebrates. Unlike most other eukaryotes that contain numerous mitochondria, trypanosomes have a single mitochondrion, which is highly elongated and reticulated. In contrast to yeast and mammals, with tens to hundreds of mitochondrial DNA (mtDNA) nucleoids distributing throughout the mitochondrial compartment, the mitochondrial genome of trypanosome is restricted to a specific structure at the basis of the flagellum known as kinetoplast (kDNA) [1]. During the trypanosome life cycle and its adaptation to different hosts and environments, the shape of its mitochondrial compartment undergoes spectacular changes that reflect its functional plasticity [2]. Indeed, the mitochondrion of this parasite exists in at least two major forms: (i) the fully active and developed one characteristic for the PCF that harbors the oxidative phosphorylation complexes (OXPHOS) for energy production [3, 4] and (ii) the functionally less active and morphologically reduced form found in the bloodstream, with energy produced exclusively through glycolysis since OXPHOS is repressed [5]. These modifications correlate with the adaptation of the parasite to the proline-rich hemolymph and tissue fluids of the blood-feeding tsetse fly and the glucose-rich blood of a mammalian host [6]. Since studies in mammals and in yeast having unraveled tight links between mitochondrial bioenergetics, morphology and dynamics [7–9], it is tempting to assume that such close relationships not only regulate mitochondrial function and energy metabolism, but also the life cycle and pathogenic potential of trypanosomes. However, in contrast to fungi and metazoa, where mitochondrial morphology and fusion/fission dynamics have been thoroughly characterized [10, 11], little is known about the morphology and dynamics of mitochondria in trypanosomes.

In most eukaryotes, mitochondria are very dynamic, alternating between two major events, fusion and fission. Mitochondrial dynamics allows to maintain mitochondrial morphology, distribution and size [8]. The main proteins involved in these events are large GTPases belonging to the dynamin superfamily protein (DSP), such as Dnm1/Drp1 in yeast and mammalian cells, which are soluble protein transiently recruited to mitochondrial and peroxisomal membranes to mediate organelle division [12, 13]. Mammalian mitochondrial fusion requires fusion of outer mitochondrial membrane (OMM) followed by fusion of inner mitochondrial membrane (IMM) which is carried out by mitofusins 1 and 2 (Mfn1 and Mfn2) and optic atrophy 1 (Opa1), respectively [14, 15]; while mitofusins are anchored to the OMM and face the cytosol, Opa1 localizes to the mitochondrial inter-membrane space (IMS). Alteration of fusion/fission dynamics and of the equilibrium between them provokes mitochondrial fragmentation or hyperfusion [12, 16], and is linked to diseases, notably neuropathies [17].

The investigation of mitochondrial fission and fusion in trypanosomes has been relatively limited until now. Available evidence indicates that mitochondrial fission is restricted to cell division [18, 19] and may not occur continuously [18, 20] but can be induced artificially [19, 21]. Evidence for mitochondrial fusion in trypanosomes came 25 years ago from *in vivo* genetic exchange of mtDNA [22]. Finally, a recent study of mitochondrial dynamics in another trypanosomatid, *Crithidia fasciculata*, unveiled the occurrence of mitochondrial fission, fusion, and sliding throughout the cell cycle, pointing to the existence of mitochondrial fusion and fission machineries in trypanosomatids [23].

Two decades of research have revealed that numerous mitochondrial fission and fusion factors that, despite significant differences, appear conserved in fungi and mammals [24]. In plants, mitochondria also fuse and divide. While plant DSPs involved in fission are known, factors directly involved in fusion have not been identified, suggesting the existence of a fusion machinery differing from that of yeast and mammals [25, 26], [27]. So far, only two dynamin-like proteins (*Tb*Dlp) have been described in *T. brucei* (Tb927.3.4720 and Tb927.3.4760): their ablation impairs mitochondrial division and endocytosis and leads to cytokinesis arrest [19, 28, 29]. *Tb*Dlp thus appears to be the functional homolog of Drp1/Dnm1 in mammals and yeast, but its fission activity on mitochondria seems restricted to the cytokinesis [19]. Another presumed DSP (*Tb*MfnL, Tb927.7.2410) has been also reported to be involved in mitochondrial fission [30]. To date, however, no protein involved in mitochondrial fusion has been identified in trypanosomes.

In this study, we revisited the function of the trypanosome DSP *Tb*MfnL [30]. Phylogenic analysis reveals that this protein defines a new family of DSP found in both eukaryotes and prokaryotes but absent from organisms containing Mfn or Opa1 homologs (fungi and mammals) and from plants. We find that excess *Tb*MfnL leads to increased mitochondrial branching and interconnexion and we show that it is anchored to the mitochondrial inner membrane, facing the matrix. Taken together, these findings suggest that *Tb*MfnL is not involved in fission, but mediates or modulates mitochondrial interconnection and branching.

## RESULTS

### 1 Sequence analysis of *Tb*MfnL identifies it as a member of a novel dynamin family

To identify potential fusion players, we started our search by careful re-examination of trypanosome genomic and proteomic databases for proteins showing sequence homology to known fusion factors (Opa1/Mfn1-2/Fzo1). We were only able to identify the previously reported dynamin-related protein, named *Tb*Mfn-Like or *Tb*MfnL (Tb927.7.2410) [30], that displays a domain organization that has more similarities with mammalian and yeast fusion factors of the outer membrane (Mfn2/Fzo1) than of the inner membrane (Opa1) (**Figure 1A and Table S1**). Analysis of the *Tb*MfnL sequence revealed several motifs and domains, including an N-terminal region containing a predicted mitochondrial targeting signal (MTS) with potential cleavage sites (**Figure S1**), followed by a GTPase domain with conserved G1 to G4 motifs and two consecutive C-terminal transmembrane domains (TMs) (**Figure 1A and Figures S1/S2**). The presence of an N-terminal MTS makes it unlikely for *Tb*MfnL to localize to the outer membrane, as Mfn/Fzo, and the absence of an N-proximal TM domain, which prevents Opa1 import into the matrix, suggests that *Tb*MfnL is imported into the matrix and does not represent a functional homolog of the fusion factors Opa1 or Mfn.

**Figure 1.**
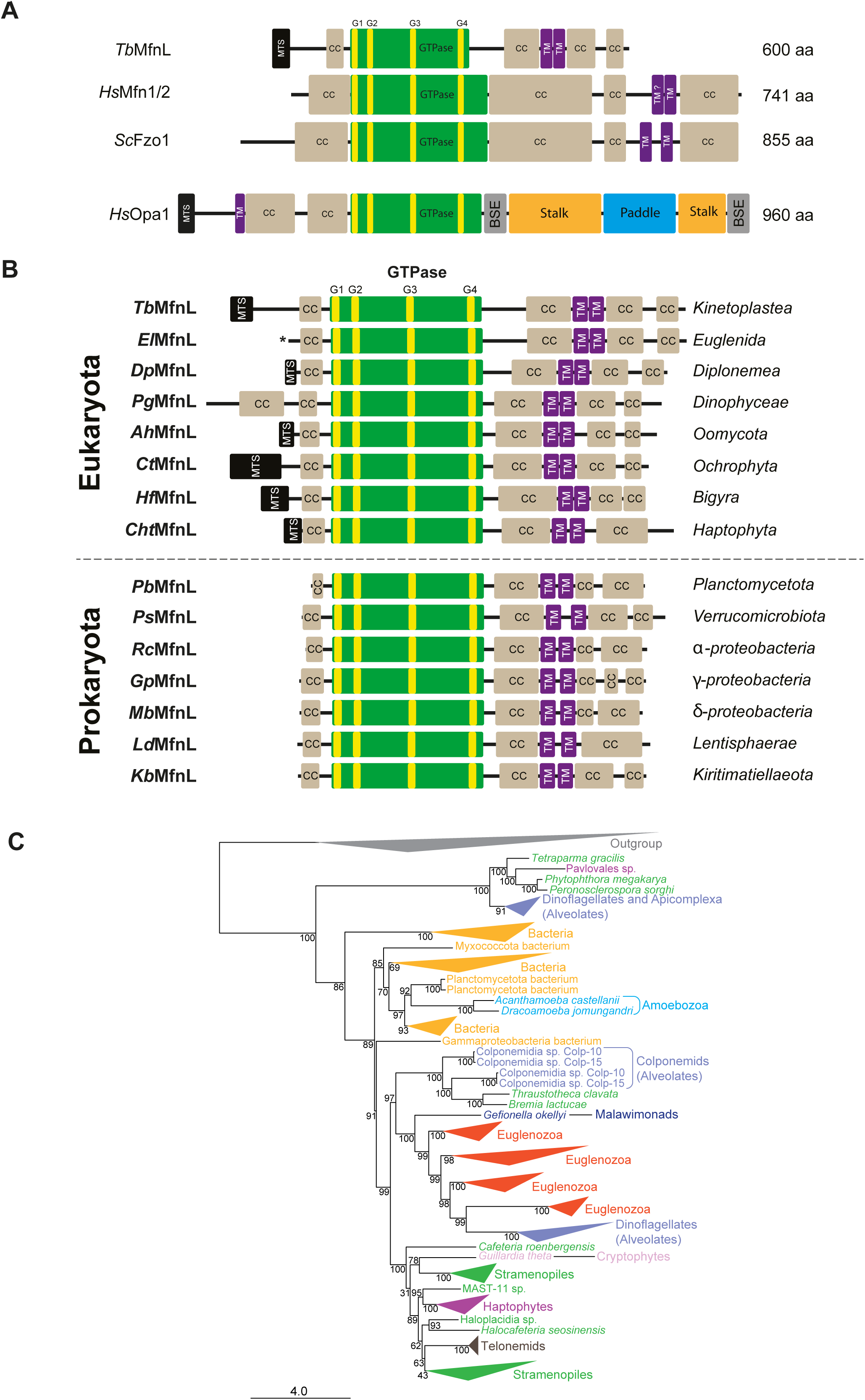
Sequences homologous to *Tb*MfnL and domain organization of the proteins. **(A)** Structure comparison between mitochondrial dynamin-superfamily proteins (DSPs): *T. brucei Tb*MfnL, human mitofusin 1 or 2 (*Hs*Mfn1/2), yeast Fzo1 (*Sc*Fzo1) and human Opa1 (*Hs*Opa1). MTS, predicted mitochondrial targeting sequence; CC, coiled coil region; green box, GTPase domain with the four GTP-binding motifs in yellow; TM, transmembrane span; BSE, bundle signaling element; Stalk domain; Paddle domain. Two TM domains were initially proposed for Mfn1/2, but recently, Mattie et al. (2018) [71] reported the presence of only a single TM domain, with the C-terminus extending into the IMS. This contrasts with Fzo1 and *Tb*MfnL, which are proposed to have a U-shaped structure. Therefore, we have indicated this as TM? in the figure. Structure predictions were performed using PSIPRED [72]. More details and sequence alignment are shown in **Figure S2**. **(B)** Domain organization of several identified proteins from various organisms. MTS, predicted mitochondrial targeting sequence; CC, coiled-coil region; green box, GTPase domain with the four GTP-binding motifs in yellow; TM, transmembrane span; Structure predictions were performed using PSIPRED. More details and sequence alignment are shown in **Figure S2**. *Tb*MfnL, *Trypanosoma brucei* (ID: Tb927.7.2410); *El*MfnL, *Euglena longa* (ID: EP00668_Euglena_longa_P029018); *Dp*MfnL, *Diplonema papillatum* (ID: KAJ9468802.1); *Pg*MfnL, *Polarella glacialis* (ID: CAE8737761.1); *Ah*MfnL, *Achlya hypogyna* (ID: OQR970001); *Ct*MfnL, *Chaetoceros tenuissimus* (ID: GFH50037.1); *Hf*MfnL, *Hondaea fermentalgiana* (ID: GBG23909.1); *Cht*MfnL, *Chrysochromulina tobinii* (ID: KOO35358.1); *Pb*MfnL, *Planctomycetaceae bacterium* (ID: MBV8881356.1); *Ps*MfnL, *Pedosphaera* sp. (ID: MCH2381658.1); *Rc*MfnL, *Rubrimonas cliftonensis* (ID: WP_093253540.1); *Gp*MfnL, *Gammaproteobacteria* bacterium (ID: RTZ59088.1); *Mb*MfnL, *Myxococcales* bacterium (ID: MCA9558847.1); *Ld*MfnL, *Lentisphaerae* bacterium (ID: NLB69313.1); *Kb*MfnL, *Kiritimatiellae* bacterium (ID: MBO7223041.1). The accession number for *Euglena longa* was obtained from Eukprot V3 web site. **(C)** Maximum likelihood phylogenetic tree of eukaryotic and prokaryotic DSPs. The tree was reconstructed using the LG+C60+G4 model of sequence evolution. Ultrafast bootstrap support values are shown on each branch. Taxa are colored according to their taxonomic affiliation. Groups of sequences belonging to the same phylum are collapsed (see **Figure S3** for the complete tree).

We next investigated whether this DSP could be conserved in other organisms. Using Blastp with *Tb*MfnL as a query to search various protein databases, we identified well-conserved sequences in relatives of *Trypanosoma* belonging to the class Kinetoplastea, including free-living bodonid species (**Figure S1 and Table S1** for examples). Surprisingly, we also discovered homologous proteins with high similarity across the entire sequence in other very diverse Euglenozoa species, including the classes Euglenida and Diplonemea, and also in various much more distantly related eukaryotic lineages, particularly within the SAR clade, including Stramenopiles and Alveolata (**Figure 1B, Figure S2 and Table S1** for example). However, no proteins with the same general architecture were identified in the Opisthokonta (the clade that includes fungi and animals) or in plants, the only sequences showing some similarity being restricted, as anticipated, to the well-conserved GTPase domain. Unexpectedly, we also identified several sequences similar to *Tb*MfnL in bacteria, mainly in the PVC group and in Proteobacteria (**Figure 1B, Figure S2 and Table S1** for example). Analysis of several of these proteins showed again a very conserved structural and domain organization along the whole sequence, with the exception of the absence of the potential mitochondrial targeting sequence (MTS) in prokaryotes (**Figure 1B and Figure S2**).

We conducted a maximum likelihood phylogenetic analysis of the identified sequences, incorporating several DSPs from mammals and yeast, as well as the cyanobacterial homolog DBLP as outgroup sequences (**Figure 1C and Figure S3**). Most eukaryotic sequences branched within a well-supported (99% ultrafast bootstrap support –UFBS-) group. Despite the fact that sequences within this clade tended to group according to their taxonomic origin, we observed several unexpected relationships (e.g., dinoflagellates branched within the Euglenozoa, and cryptophytes and haptophytes within the Stramenopiles). Moreover, colponemids (a group of deep-branching alveolates) formed a fully-supported (100% UFBS) group with two stramenopile sequences. Bacterial sequences formed a paraphyletic group at the base of this main eukaryotic clade, with two ameobozoan species (*Acanthamoeba castellanii* and *Dracoamoeba jomungandri*) robustly branching (97% UFBS) within one of the bacterial subgroups. This unexpected relationship strongly suggested that these amoebae acquired this gene by horizontal gene transfer (HGT) from bacterial donors. By contrast, the anomalous relationships among eukaryotic taxa observed within the main eukaryotic clade were more difficult to explain, although multiple gene duplications and losses appeared to have played a role to shape the current distribution of this gene in eukaryotes. The class Diplonemea provided evidence for this hypothesis, with two highly supported (100% UFBS) groups of sequences of the same genera (*Diplonema*, *Lacrimia*, and *Rhynchopus*) branching paraphyletically at the base of a large clade of dinoflagellate (Alveolata) sequences (**Figure S3**). A small group of sequences containing stramenopile, alveolate, and haptophyte species branched sister to all the other prokaryotic and eukaryotic sequences and most likely represented an ancient paralog of this gene, giving additional evidence for the role of duplication in the evolution of this gene. Furthermore, it is very likely that HGT between different eukaryotic groups also occurred, as in the case of the dinoflagellate sequences cited above, which appear to have acquired this gene from euglenozoan donors, and in the case of the haptophyte and cryptophytes sequences that branched within the large group of stramenopile sequences. Given the specific architecture of this DSP, its large phylogenetic distance to other previously known protein families, and its wide and distinct taxonomic representation in prokaryotes and eukaryotes –but its absence in animals and plants-strongly indicate that it represents a novel DSP associated with membrane dynamics.

### 2 *Tb*MfnL is localized in the mitochondrion

To determine the subcellular localization of *Tb*MfnL (Tb927.7.2410), we added a 10xHA tag at the C-terminal extremity of the endogenous *Tb*MfnL (*Tb*MfnL_::10HA_) in both PCF and BSF trypanosomes (**Figure 2A**). C-terminal tagging was used to avoid disruption of the potential N-terminal MTS. Western blot analysis with anti-HA antibody showed that *Tb*MfnL_::10HA_ is about 2-fold more expressed in PCF than BSF, which is in agreement with the data obtained through SILAC and transcriptomic analysis for this protein [31, 32] (**Figure 2B**). The localization of the *Tb*MfnL protein was studied in PCF by immunofluorescence using threonine dehydrogenase (Tdh, [33]) as a mitochondrial matrix marker (**Figure 2C**). *Tb*MfnL_::10HA_ and Tdh clearly colocalized in the mitochondrial compartment, but *Tb*MfnL depicted a less homogeneous mitochondrial distribution (**Figure 2C, Merge**). Unfortunately, we could not use the anti-Tdh immune serum for immunofluorescence analysis on BSF. Consequently, we performed a labeling of the mitochondria, with an anti-HSP60 (mitochondrial maker). Like observed in PCF, *Tb*MfnL_::10HA_ and HSP60 colocalized in the mitochondrial compartment (**Figure 2C, Merge)**. The presence of a N-terminal targeting sequence and its localization by fluorescence microscopy demonstrated that *Tb*MfnL is a mitochondrial protein. These data are consistent with the mitochondrial localization of the C-terminal GFP-tagged *Tb*MfnL reported by the TrypTag database (cellular localization of trypanosome proteins [34]). Of note, the N-terminal GFP-Tagged *Tb*MfnL exhibited a cytoplasmic localization in TrypTag database [34], likely due to masking of the MTS pre-sequence.

**Figure 2.**
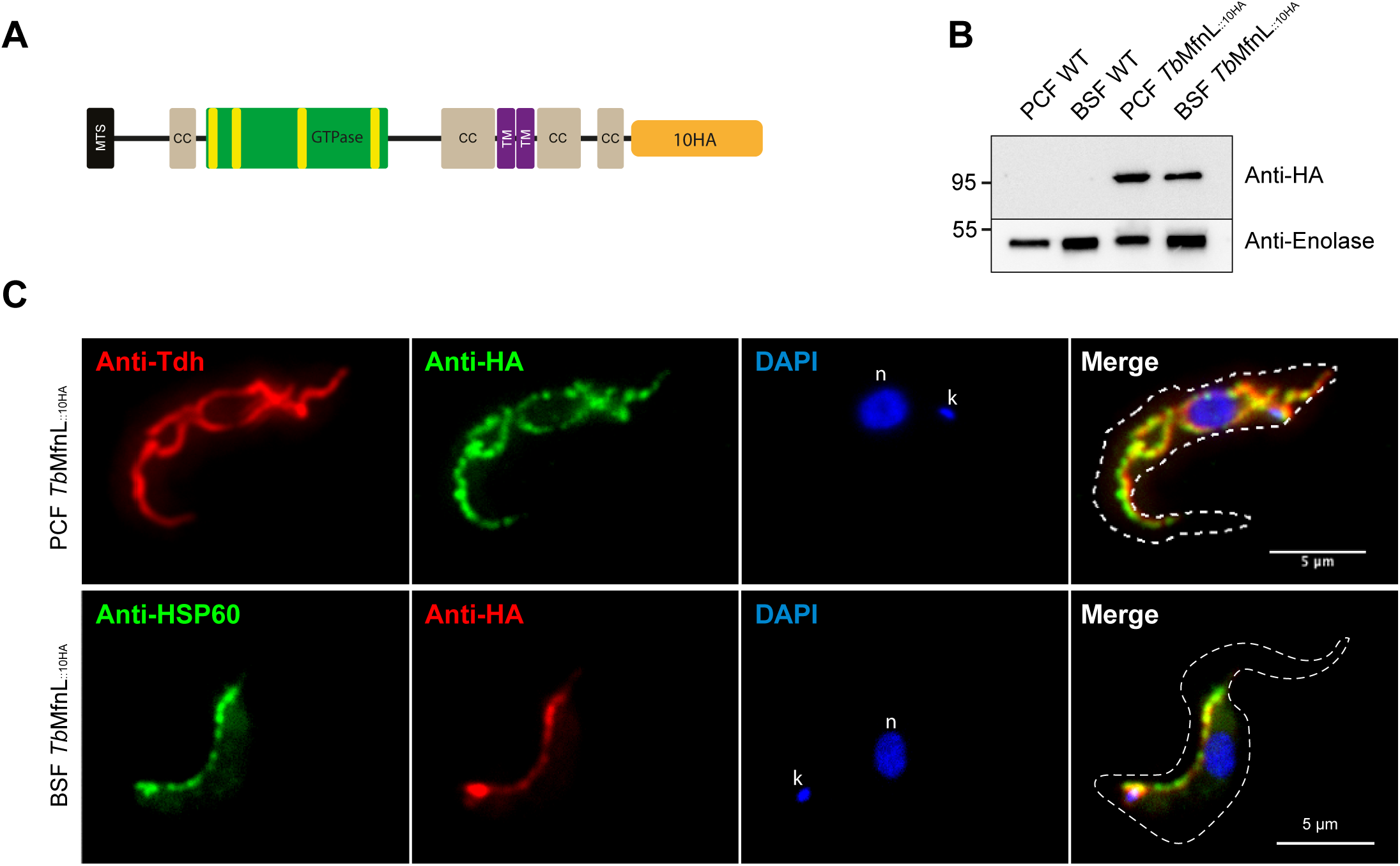
Subcellular localization of *Tb*MfnL. **(A)** Schematic representation of *Tb*MfnL endogenously tagged at its C-terminus extremity with 10xHA. Only one of the two alleles is tagged. **(B)** Western blotting of whole-cell extracts (5×10^6^ cells) of *T. brucei* PCF or BSF wild-type cells and PCF or BSF cells expressing *Tb*MfnL_::10HA_ revealed with an anti-HA antibody. It should be noted that Enolase used as a loading control is about 3 to 4 times more expressed in BSF than in PCF according to Hannaert *et al*. 2003 [73]. **(C)** Subcellular localization of *Tb*MfnL_::10HA_ in PCF and BSF. Colocalization of *Tb*MfnL_::10HA_ (anti-HA antibody) with matrix mitochondrial threonine dehydrogenase for PCF (Anti-Tdh) and mitochondrial HSP60 for BSF (Anti-HSP60) analyzed by standard immunofluorescence. n, nucleus; k, kinetoplast. The scale bar represents 5 µm.

### 3 Inactivation and silencing of *Tb*MfnL does not alter mitochondrial shape

Live microscopy of yeasts and cultured mammalian cells has revealed that mitochondrial morphology is continuously remodeled by antagonistic fission and fusion reactions [16, 35]. Under normal conditions, the fusion/fission balance is established in favor of fusion and the mitochondria appear filamentous. Inhibition of fusion leads to mitochondrial fragmentation by ongoing fission [16] and reversely enhancing fusion leads to hyperfused mitochondria with increased branching and interconnection [12].

To investigate the role of *Tb*MfnL, we first inactivated the *Tb*MfnL gene by CRISPR/Cas9 [36]. Both *Tb*MfnL alleles were inactivated by integration of a puromycin resistance marker cassette into both PCF and BSF cloned cells (**Figure 3A**). Cloned parasites were analyzed by PCR to select cell lines carrying puromycin resistance gene insertions in both *Tb*MfnL alleles (*Tb*MfnL^−/-^) (**Figure 3B**). No significant impact on cell growth was observed in these PCF and BSF null-mutants over a period of 7 days compared to parental cells (**Figure S4**) and no significant difference in the structure of the *Tb*MfnL^−/-^ mitochondrion was observed by staining mitochondrion with rhodamine-123 or immunofluorescence analyses with the anti-HSP60 immune serum, in both PCF and BSF mutant cell lines, respectively (**Figure 3C and 3D**). To confirm that the CRISPR/Cas9 strategy leads to *Tb*MfnL gene inactivation, we repeated this process in a strain endotagged with a C-terminal 3xTy1 tag in order to follow the expression of the protein by western blot (**Figure S5A-D**) upon insertion of the puromycin resistance gene as above (**Figure S5E-G**) and deletion of part of the *Tb*MfnL gene (**Figure S5H-J**), as above. In both cases, the protein was not detected in the inactivated clones and no modification of the mitochondrial structure was observed (**Figure S5G/J)**.

**Figure 3.**
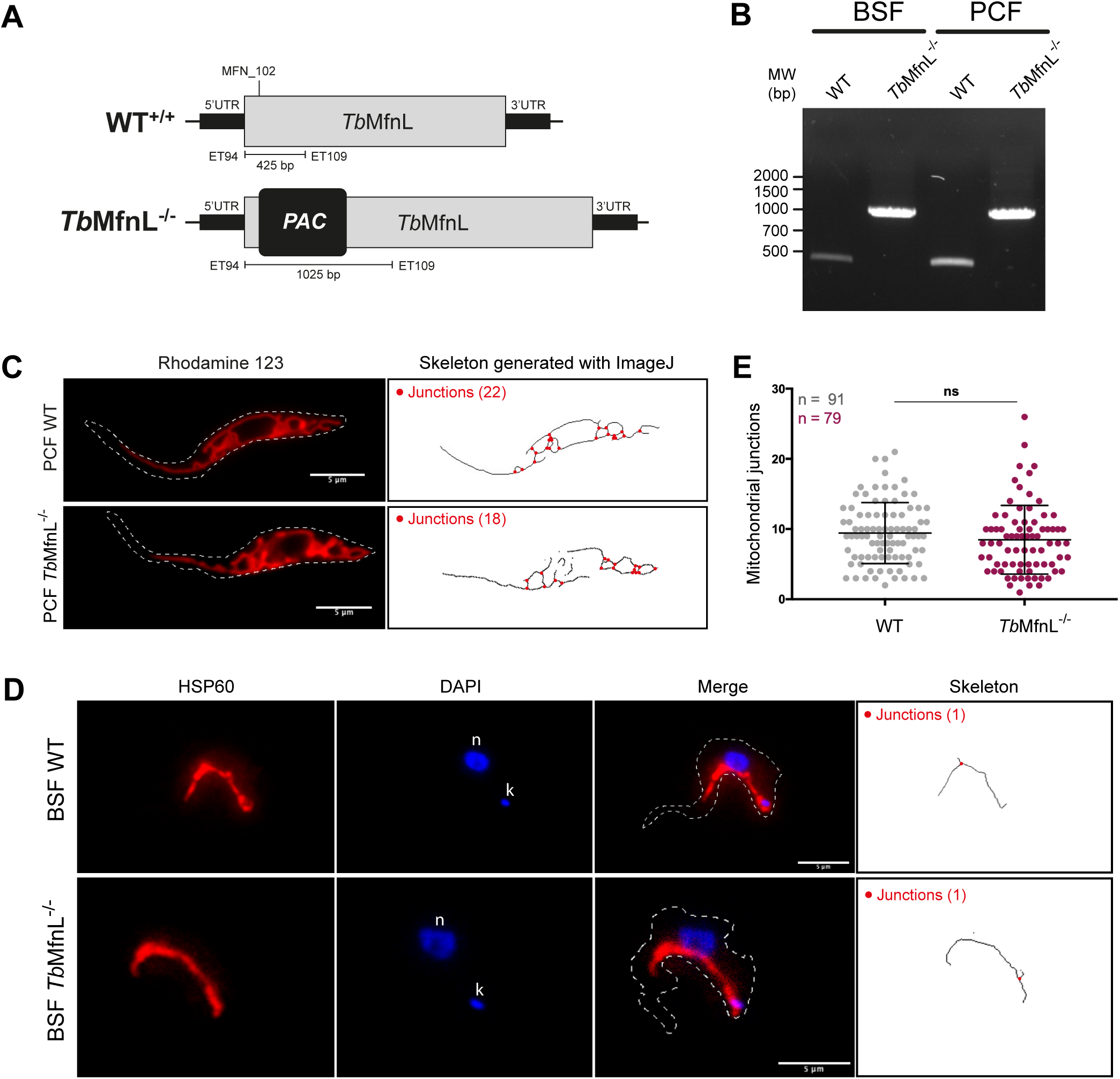
Inactivation of the *Tb*MfnL gene. **(A)** Schematic representation of *Tb*MfnL inactivation by insertion of the puromycine resistant marker (PAC). **(B)** PCR confirmation of *Tb*MfnL gene inactivation on both alleles in BSF and PCF. **(C)** The mitochondrial shape of living parental (WT) and *Tb*MfnL^−/-^ PCF cells (right panel) were designed by an ImageJ macro (Mitochondrial Junctions) from rhodamine 123 staining (left panel). **(D)** Antibody directed against HSP60 was used to label and visualized mitochondrial shape in BSF cells. Top panel: BSF parental cell (WT); bottom panel: BSF *Tb*MfnL^−/-^ cell. n, nucleus; k, kinetoplast. **(E)** Corresponding mitochondrial junctions’ quantification. Small red circles in the skeleton image indicate the junctions where three tubules converge. The scale bar represents 5 µm. Statistic: t test - Confidence interval 95% - P-value style: 0.1234 (ns); 0.0332 (*); 0.0021 (**); 0.0002 (***); <0.0001 (****).

In order to detect and quantify discrete modifications of the mitochondrial shape that may escape detection upon qualitative visual analysis, we developed an ImageJ macro (**Data S1**) for automatic quantification of mitochondrial junctions from fluorescence images (**Figure 3C/D/E**). This allowed us to establish that the number of mitochondrial junctions is equivalent in *Tb*MfnL^−/-^ and parental PCF cells (8.5±4.9 vs 9.4±4.3) confirming the qualitative analysis of the microscopy images (**Figure 3C**). The mitochondrial and cristae structures were also examined by transmission electron microscopy (TEM) in *Tb*MfnL^−/-^ PCF cells cultured with either proline or glucose as the sole carbon source. Under these conditions, no significant alterations in mitochondrial or cristae structure were observed, nor were there any noticeable effects on the growth of the mutant cells (**Figure S6**).

These findings regarding the consequences of *Tb*MfnL inactivation in PCF and BSF are not in line with a previous report describing a highly fenestrated mitochondrial network consecutive to a partial but significant inhibition (70%) of *Tb*MfnL expression by RNAi in BSF [30]. To exclude that the divergence arose from the different inactivation and silencing strategies, we conditionally down-regulated expression of *Tb*MfnL using a stem-loop RNAi construct. This approach led to a significant reduction (∼70%) of the expression of endogenously tagged *Tb*MfnL_::10HA_ in PCF (as observed in BSF [30]), but this was not paralleled by a modification of mitochondrial shape (**Figure S7A-C**). Since mitochondrial fenestration upon *Tb*MfnL silencing was observed in BSF [30], we decided to repeat the experiment using the same strain (*T. brucei* 427 BSF), the same RNAi system (p2T7-177) and the same sequence used for down-regulation [30]. As previously reported [30], we observed a significant reduction of *T*bMfnL expression (∼80% after 4 days) of the endogenously tagged *Tb*MfnL_::3HA_ in BSF (**Figure S7D-F**) that was not accompanied by mitochondrial shape modifications (**Figure S7G**). In our hands, neither inactivation nor silencing of *Tb*MfnL alters mitochondrial morphology or induces a fenestration phenotype.

### 4 *Tb*MfnL overexpression increases mitochondrial branching

We also investigated the possible role of *Tb*MfnL by conditionally overexpressing a C-terminally-tagged (3xTy1) *Tb*MfnL in PCF (^oe^*Tb*MfnL_::3Ty1_, **Figure 4A**). Expression was confirmed by western blot analysis after 3 to 5 days of tetracycline induction (**Figure 4B**). Rhodamine-123 staining showed that the parental and non-induced ^oe^*Tb*MfnL_::3Ty1_ cell lines harbor the expected reticulated mitochondrial structure, while the mitochondrion appears significantly more reticulated and branched in the tetracycline-induced ^oe^*Tb*MfnL_::3Ty1_ cells (**Figure 4C, Figure S8**). After 5 days of induction, the average number of mitochondrial junctions, quantified with the ImageJ macro, appeared more than 2 times higher in induced ^oe^*Tb*MfnL_::3Ty1_ cells compared to non-induced or parental cells (**Figure 4D**). In order to estimate the level of *Tb*MfnL overexpression achieved in *T. brucei* PCF, we overexpressed *Tb*MfnL_::3Ty1_ in the *Tb*MfnL_::3Ty1_ background, in which the endogenous *Tb*MfnL alleles were 3Ty1-tagged (clone 2A11; **Figure S5A-D**). Western blot analysis with the anti-Ty1 antibodies revealed that expression of *Tb*MfnL_::3Ty1_ is 5.5-times increased in the resulting tetracycline-induced *Tb*MfnL_::3Ty1_/^oe^*Tb*MfnL_::3Ty1_ compared to the parental *Tb*MfnL_::3Ty1_ clone, which only expressed the endogenously tagged *Tb*MfnL_::3Ty1_ (**Figure S9A**). Overexpression of *Tb*MfnL_::3Ty1_ once again leads to alterations in mitochondrial morphology (**Figure S9B**). It is also noteworthy that the expression of *Tb*MfnL without the C-terminal 3Ty1 tag induced the same phenotype as the tagged *Tb*MfnL, confirming that the phenotype is not due to the use of a C-terminal tag (**Figure S9A/B**). These data clearly demonstrated that overexpression of *Tb*MfnL stimulates inter-mitochondrial connections.

**Figure 4.**
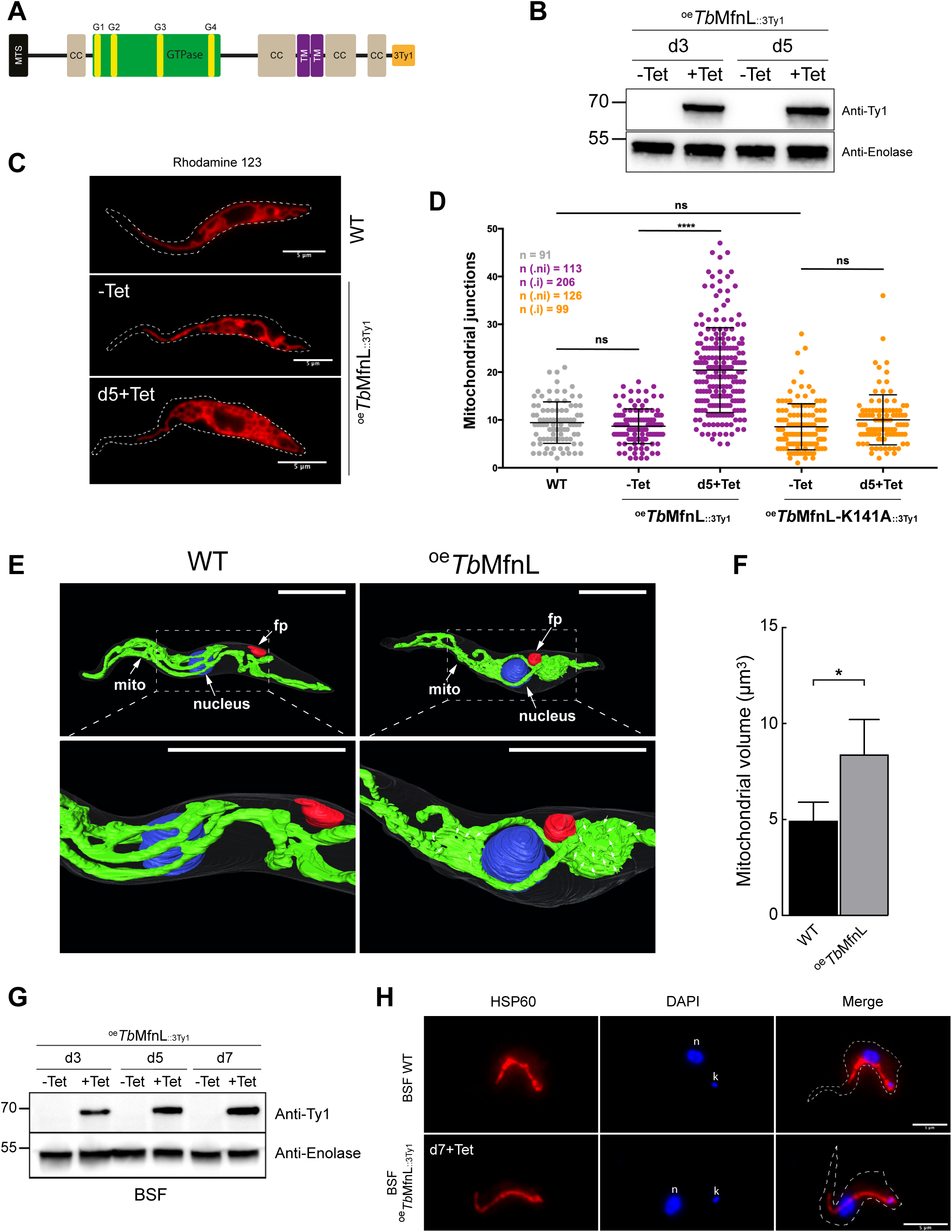
Overexpression of *Tb*MfnL and GTPase mutant. **(A)** Schematic representation of the overexpressing *Tb*MfnL protein tagged with 3Ty1 peptide at the C-terminus of the protein (^oe^*Tb*MfnL_::3Ty1_). **(B)** Western blotting of whole-cell extracts of *T. brucei* PCF cells overexpressing *Tb*MfnL_::3Ty1_. Non-induced (-Tet) and induced (+Tet) cells after 3 to 5 days (d3 and d5) were revealed with an anti-Ty1 antibody. Antibody directed against Enolase protein was used as loading control. **(C)** Mitochondrial structure analysis using rhodamine 123 staining on living cells; mitochondrial structure of PCF parental (WT) and PCF ^oe^*Tb*MfnL_::3Ty1_ cells before (- Tet) and 5 days after (+Tet) tetracycline induction. **(D)** Quantification of mitochondrial junctions’ number using Rhodamine 123 microscopy images. Mitochondrial junctions number: WT, 9.4 ± 4.3; -Tet ^oe^*Tb*MfnL_::3Ty1_ 8.7 ± 3.6; +Tet ^oe^*Tb*MfnL_::3Ty1_ 20.4 ± 8.9; -Tet ^oe^*Tb*MfnL-K141A_::3Ty1_ 8.6 ± 4.8 and +Tet ^oe^*Tb*MfnL-K141A_::3Ty1_ 10 ± 5.2. **(E)** Surface rendering of segmented PCF using SBF-SEM. Surface rendering of segmented PCF WT and ^oe^*Tb*MfnL cells, with ^e^*Tb*MfnL induced by tetracycline for 5 days, showing the cell body (transparent), nucleus (blue), mitochondrion (green), and flagellar pocket (red). 5 WT and 4 ^oe^*Tb*MfnL cells were analyzed. A magnified view of the upper images, enclosed in a box, was shown. Small white arrows mark the mitochondria-free areas within the mitochondrial network. **(F)** Determination of mitochondrial volume (µm^3^) in both WT and ^oe^*Tb*MfnL cells; WT 4.9±1 and ^oe^*Tb*MfnL 8.3±1.9. Statistic: t test - Confidence interval 95% - P-value style: 0.1234 (ns); 0.0332 (*); 0.0021 (**); 0.0002 (***); <0.0001 (****). **(G)** Western blotting of whole-cell extracts of *T. brucei* BSF overexpressing *Tb*MfnL_::3Ty1_ cells. Non-induced (-Tet) and induced (+Tet) cells after 3 to 7 days (d3, d5 and d7) were revealed with an anti-Ty1 antibody. Antibody directed against Enolase protein was used as loading control. **(H)** Subcellular localization of ^oe^*Tb*MfnL_::3Ty1_ in BSF by immunofluorescence. Antibody directed against HSP60 was used to label and visualized mitochondrial shape in parental (WT) and *Tb*MfnL_::3Ty1_ overexpressing cells. n, nucleus; k, kinetoplast. The scale bar represents 5 µm.

To characterize further inter-mitochondrial branching with higher resolution we used serial block face scanning electron microscopy (SBF-SEM), an approach allowing to characterize the overall structure of membranes and organelles within the entire cell volume [18]. With this approach, hundreds of images are collected to perform a 3D reconstruction of the whole cell, as well as spatial organization of the individual organelles within the cell. Four to five parental and ^oe^*Tb*MfnL_::3Ty1_ PCF cells were reconstructed in 3D, with a particular emphasis on the mitochondrion, the nucleus, the flagellar pocket and the cell outline. We have not been able to correctly identify the kinetoplast DNA on the electron microscopy images. However, since kDNA is in close contact with the flagellar pocket, we have assumed its position. Other organelles (glycosomes, acidocalcisomes, Golgi, flagellum, etc.) are not represented either in order to not overload the 3D representation. As expected, the parental PCF cells showed a reticulated mitochondrion along the whole cell (**Figure 4E**), a rather central nucleus and the flagellar pocket on the posterior side. The length of the parental and ^oe^*Tb*MfnL_::3Ty1_ cells, varied between 20.4±3.3 and 23±4.7 µm, respectively. As observed by fluorescence microscopy, ^oe^*Tb*MfnL_::3Ty1_ mitochondrion showed more reticulation (**Figure 4E**). Unexpectedly, in some cells this reticulation was even more prominent at the edges of the flagellar pocket, where the kDNA is located, as illustrated in **Figure 4E**, an observation that we had not identified by immunofluorescence microscopy approaches. This hyper-reticulated structure (leaving a few gaps in the mitochondrion) formed a globular structure and probably encapsulated the kDNA (**Figure 4E**). This structure called kDNA pocket has been already described in BSF where kDNA replicates [20], but is even more prominent in the ^oe^*Tb*MfnL_::3Ty1_ cells. We determined that the mitochondrial volume in PCF cells (4.9 ± 1 µm³, **Figure 4F**) aligns with previous reports (2.5–3 µm³, [2]) and as expected, is nearly twice that of BSF cells (1.1–3 µm³, [18, 20]). As anticipated, the mitochondrial volume in *^oe^TbMfnL_::3Ty1_* cells is nearly doubled compared to parental cells (8.3 ± 1.9 µm³, **Figure 4F**). Additionally, TEM analysis of the mitochondrial and cristae structures in ^oe^*Tb*MfnL_::3Ty1_ cells cultured with either proline or glucose as the sole carbon source revealed no significant alterations (**Figure S6**).

We also investigated the effect of *Tb*MfnL expression in the BSF trypanosomes, where the mitochondrial compartment is smaller and composed of a shorter and poorly interconnected mitochondrion. Expression of ^oe^*Tb*MfnL_::3Ty1_ was confirmed by western blot (**Figure 4G**), but unlike in PCF, the mitochondrial structure (labeled here with mitochondrial HSP60) was not altered, remaining tubular like in the parental cells (**Figure 4H**). Induction over a longer period (up to 21 days) did not show mitochondrial alterations.

### 5 *Tb*MfnL overexpression increases mitochondrial branching in a GTPase dependent manner

All dynamin-like proteins possess a GTP-binding domain with highly conserved G1-G4 motifs. They can be inactivated by mutations in the G1 motif that lower the efficiency of GTP-binding and hydrolysis. To determine whether *Tb*MfnL activity requires a functional GTPase domain, the key lysine 141 in the highly conserved G1 motif was converted to alanine (K141A), according to previously characterized GTP-binding domains from other organisms [37–39]. The K141A mutation was introduced into the *Tb*MfnL_::3Ty1_ sequence, which was overexpressed in the PCF ^oe^*Tb*MfnL-K141A_::3Ty1_ (**Figure 5A**) and *Tb*MfnL_::3Ty1_ PCF backgrounds (*Tb*MfnL_::3Ty1_/^oe^*Tb*MfnL-K141A_::3Ty1_, **Figure S9**). Expression of *Tb*MfnL-K141A_::3Ty1_ was confirmed by western blot analyses after 3 to 5 days of tetracycline induction in the ^oe^*Tb*MfnL-K141A_::3Ty1_ cell line (**Figure 5B**) and its overexpression was estimated as 3-fold in the *Tb*MfnL_::3Ty1_/^oe^*Tb*MfnL-K141A_::3Ty1_ mutant compared to the parental *Tb*MfnL_::3Ty1_ clone (**Figure S9A**). *Tb*MfnL-K141A_::3Ty1_ localized to the mitochondria (**Figure 5C**), as expected, but did not alter mitochondrial shape (**Figure 5C/D, Figure S8**) nor the number of mitochondrial junctions (**Figure 4D**). To exclude the possibility that concentration-dependent effects (due to the twofold lower expression of *Tb*MfnL-K141A_::3Ty1_ compared to the parental *Tb*MfnL_::3Ty1_) might prevent K141A from inducing membrane branching, rather than its inactivated GTPase activity, we equalized the expression levels of wild-type *Tb*MfnL_::3Ty1_ and mutant *Tb*MfnL-K141A_::3Ty1_. The tetracycline concentration used for *Tb*MfnL_::3Ty1_ expression was reduced from 1 µg/ml to 0.05 µg/ml. Under these conditions, both variants displayed comparable expression levels at 3 and 5 days (**Figure S10A**); however, hyper-reticulation was observed exclusively in cells overexpressing *Tb*MfnL_::3Ty1_ (**Figure S10B**). This clearly demonstrates that *Tb*MfnL is a dynamin-related protein that relies on a functional GTPase domain to modulate mitochondrial branching in trypanosomes.

**Figure 5.**
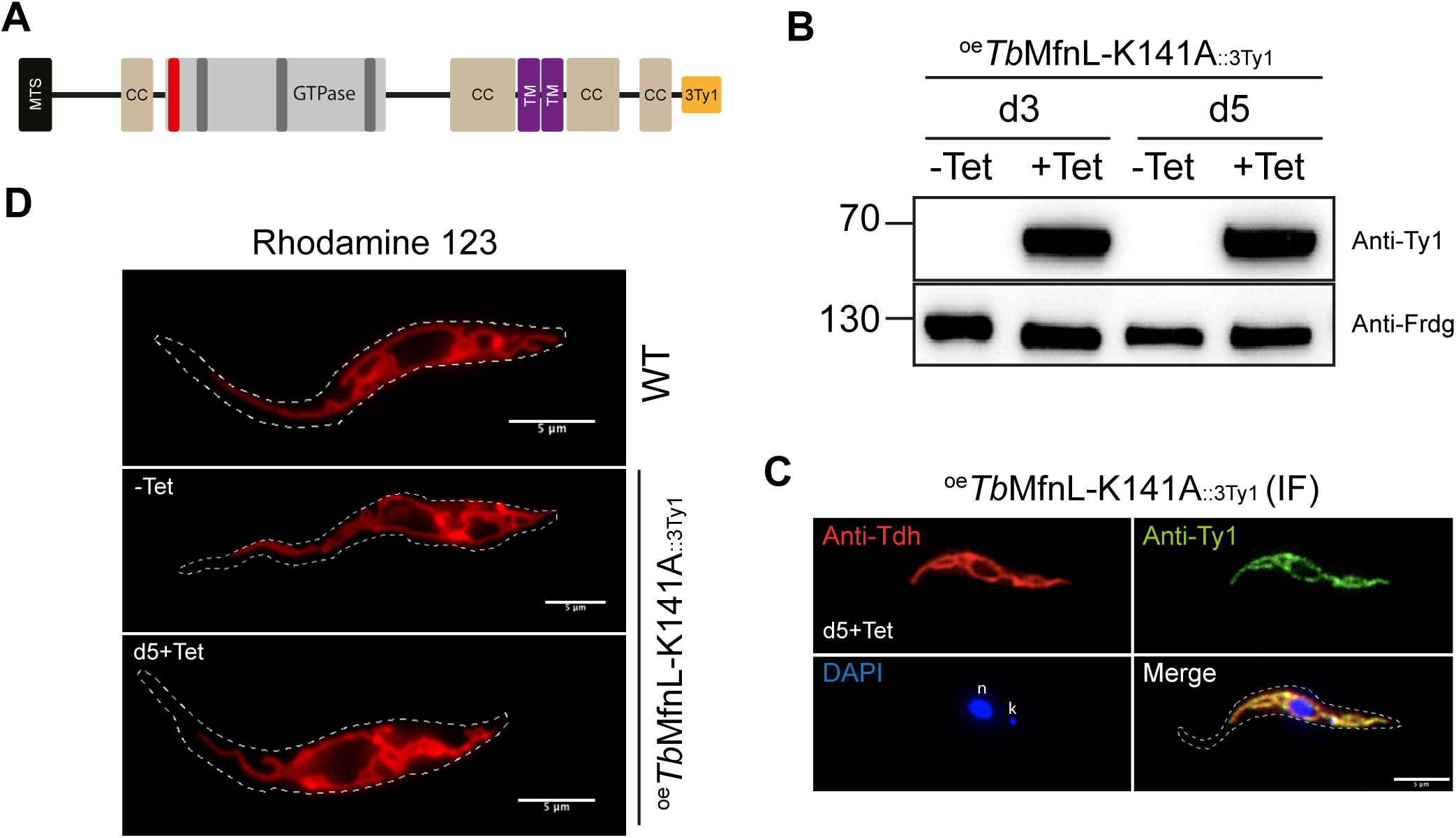
Overexpression of GTPase mutant(A) Schematic representation of the overexpressing *Tb*MfnL-K141A_::3Ty1_ mutant protein tagged with 3Ty1 peptide at the C-terminus of the protein (^oe^*Tb*MfnL-K141A_::3Ty1_). **(B)** Western blotting of whole-cell extracts of *T. brucei* PCF overexpressing *Tb*MfnL-K141A_::3Ty1_ cells; (-Tet) non-induced and (+Tet) induced cells for 3 or 5 days (d3 and d5). Antibody directed against glycosomal Fumarate reductase protein (Frdg) was used as loading control. **(C)** Submitochondrial localization of ^oe^*Tb*MfnL-K141A_::3Ty1_ in PCF by immunofluorescence; Colocalization of ^oe^*Tb*MfnL-K141A_::3Ty1_ (anti-Ty1) with matrix mitochondrial Threonine dehydrogenase (Anti-Tdh) was analyzed by standard immunofluorescence, after 5 days induction (d5+Tet). n, nucleus; k, kinetoplast. **(D)** Mitochondrial structure analysis using rhodamine 123 staining on living cells; mitochondrial structure of PCF parental (WT) and PCF ^oe^*Tb*MfnL-K141A_::3Ty1_ cells before (-Tet) and 5 days after (+Tet) tetracycline induction. The scale bar represents 5 µm.

The mitochondrial localization of ^oe^*Tb*MfnL_::3Ty1_ and ^oe^*Tb*MfnL-K141A_::3Ty1_ was also confirmed by electron microscopy and immunogold-labeling using an anti-Ty1 antibody. After induction and immunogold-labeling, both proteins are clearly localized in the mitochondrion (**Figure S11A/B-c/d**) as compared to controls with non-induced cells (**Figure S11A/B-a/b**), and confirmed by gold-particles quantification (**Figure S11C**).

### 6 Submitochondrial localization of *Tb*MfnL

Immunofluorescence analyses showed that ^oe^*Tb*MfnL_::3Ty1_ co-localized with Tdh (**Figure 6A**), as previously described for the endogenously-tagged *Tb*MfnL_::10HA_ (**Figure 2C**). To increase the resolution of *Tb*MfnL localization we performed Ultrastructure Expansion Microscopy (U-ExM), which increases the size of a sample while preserving its ultrastructure [40–42]. Using this approach, we showed that both ^oe^*Tb*MfnL_::3Ty1_ (**Figure 6B**) and the endogenous *Tb*MfnL_::10HA_ (**Figure 6C**) displayed mitochondrial localization with punctate, non-homogeneous distribution patterns, which differed from the localization of the matrix marker (Tdh). These findings suggest an association of the *Tb*MfnL protein with mitochondrial membranes.

**Figure 6.**
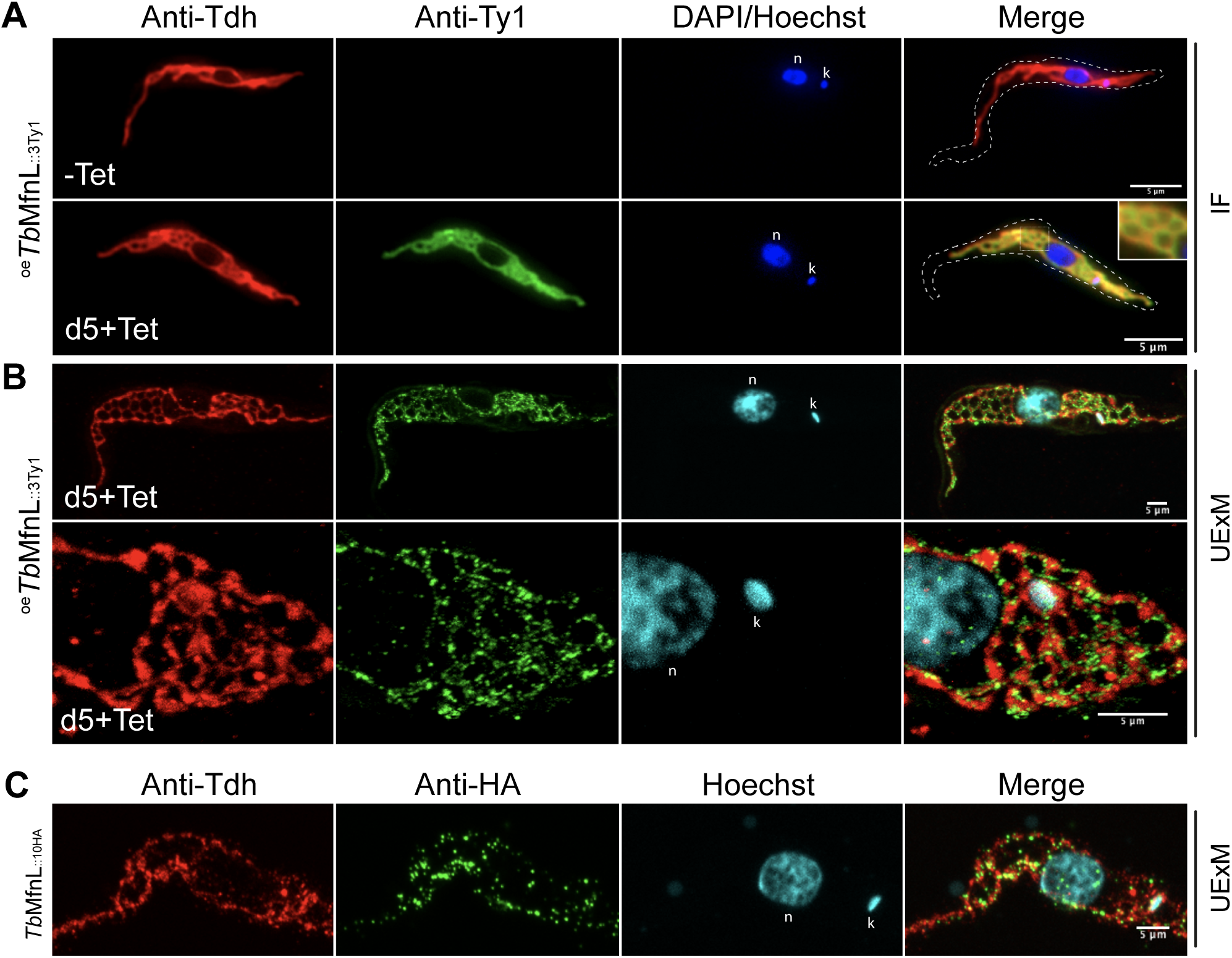
Submitochondrial localization of *Tb*MfnL. Subcellular localization of ^oe^*Tb*MfnL_::3Ty1_ in PCF by immunofluorescence and Ultra Expansion Microscopy (UExM); Colocalization of ^oe^*Tb*MfnL_::3Ty1_ (anti-Ty1) with matrix mitochondrial Threonine dehydrogenase (Anti-Tdh) was analyzed by standard immunofluorescence **(A)** and by UExM with an expansion factor of ∼4.4 **(B)**, after 5 days induction (d5+Tet). **(C)** Subcellular localization of ^oe^*Tb*MfnL_::10HA_ in PCF analyzed by UExM. n, nucleus; k, kinetoplast. The scale bar represents 5 µm.

### 7 Mitochondrial targeting and membrane anchoring of *Tb*MfnL

To investigate the mechanisms ensuring mitochondrial targeting of *Tb*MfnL, we first addressed the two domains that may determine *Tb*MfnL localization: (i) the N-terminal MTS which may facilitate targeting to the inner membrane, intermembrane space and matrix localization, and (ii) the C-proximal/terminal TMs which could mediate membrane anchoring. To investigate their capacity to target proteins to mitochondria, we fused the MTS domain, or the two transmembrane domains of *Tb*MfnL, to GFP. Fusion of the MTS at the N-terminus of GFP (_MTS::_GFP) fully targeted GFP to the mitochondria (**Figure 7A**). Interestingly, western blot analyses showed that _MTS::_GFP and cGFP (cytosolic GFP) have the same apparent size (**Figure 7B**), suggesting that the predicted MTS (**Figure S1**) was indeed cleaved by matrix processing peptidase upon import of _MTS::_GFP across the inner membrane. In the same line, ^oe^*Tb*MfnL_::3Ty1_ and ^oe^*Tb*MfnL-K141A_::3Ty1_ showed the same apparent size as ^oe^*Tb*MfnL-ΔMTS_::3Ty1_ by western blot analyses, suggesting that the MTS is also cleaved in its natural context (**Figure 7C**). In contrast, western blot analyses showed that addition of the two C-proximal TMs at the C-terminal extremity of GFP (GFP_::TM_) induces a size shift (**Figure 7B**). Immunofluorescence analyses revealed that GFP_::TM_ was not targeted to the mitochondrion or cytosol, but instead to others distinct structures (**Figure 7A**). However, it is important to note that this localization could simply represent unspecific membrane-targeting by hydrophobic TMs. This inability of *Tb*MfnL TMs to mediate mitochondrial targeting indicates that they differ from the TMs of other mitofusins, which are responsible for mitochondrial outer membrane localization [43].

**Figure 7.**
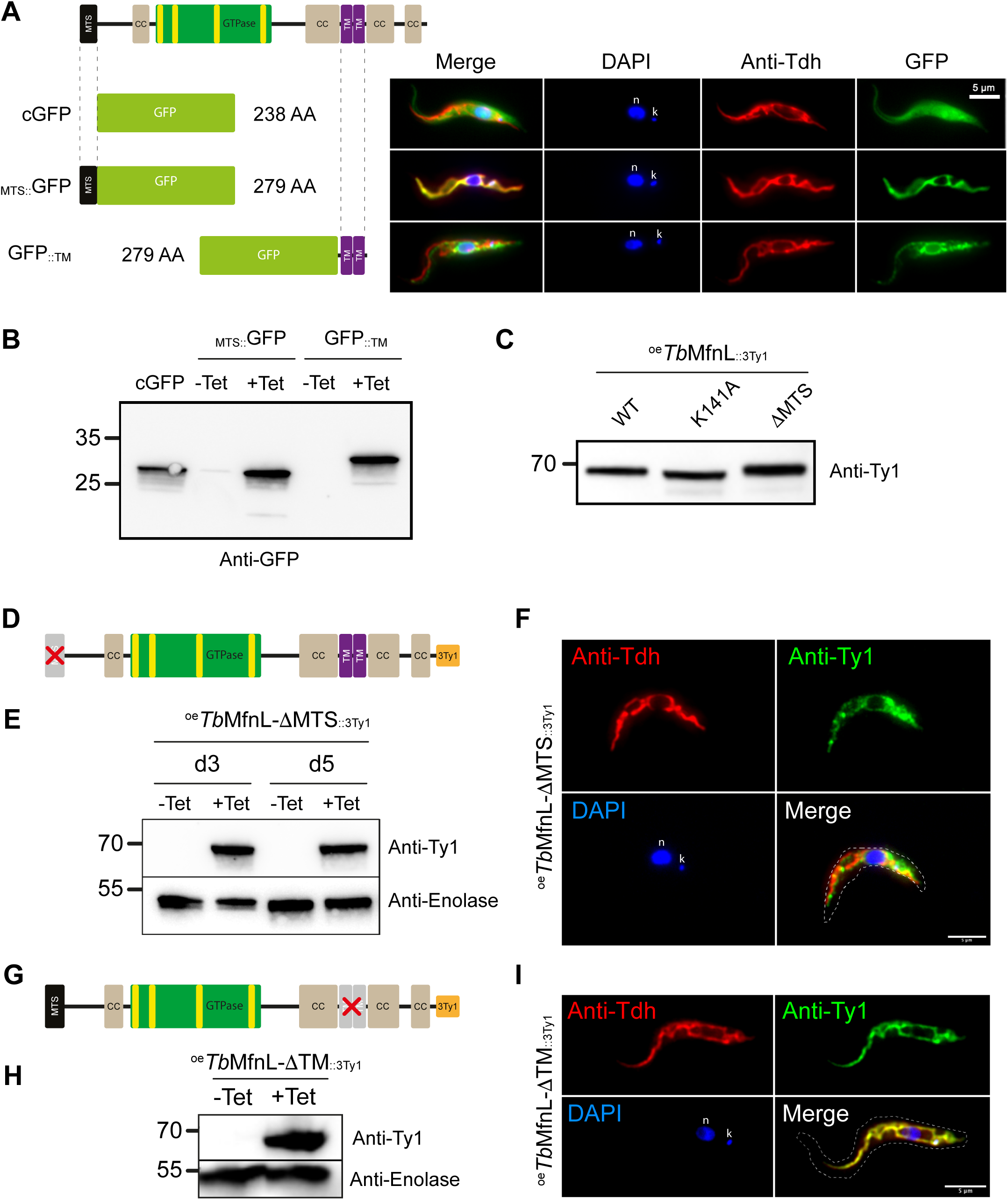
Role of *Tb*MfnL domains. **(A)** Localization of the GFP according to the different tags fused. Colocalization of GFP with the mitochondrial matrix marker threonine dehydrogenase (Anti-Tdh) was assessed by standard immunofluorescence after 5 days of tetracycline induction for MTS_::GFP_ and GFP_::TM_. The “c” in cGFP stands for constitutive expression, resulting in the cytosolic localization of GFP. MTS, mitochondrial targeting signal; TM, transmembrane domains. **(B)** Western blot analysis of GFP expression. cGFP, constitutive and cytosolic GFP. Non-induced (-Tet) and induced (+Tet) cells after 5 days. **(C)** *Tb*MfnL size analysis by western blot using anti-Ty1 antibody. **(D)** Schematic representation of N-terminus truncated *Tb*MfnL protein (first 41 amino acids) tagged with 3Ty1 peptide at the C-terminus of the protein (^oe^*Tb*MfnL-ΔMTS_::3Ty1_). **(E)** Western blotting of whole-cell extracts of *T. brucei* PCF overexpressing *Tb*MfnL-ΔMTS_::3Ty1_ cells; (-Tet) non-induced and (+Tet) induced cells for 3 or 5 days. **(F)** Subcellular localization of ^oe^*Tb*MfnL-ΔMTS_::3Ty1_ in PCF after 5 days induction. Colocalization of ^oe^*Tb*MfnL-ΔMTS_::3Ty1_ (anti-Ty1) with matrix mitochondrial Threonine dehydrogenase (Anti-Tdh) analyzed by standard immunofluorescence. **(G)** Schematic representation of truncated transmembrane domains of *Tb*MfnL protein (45 amino acids) tagged with 3Ty1 peptide at the C-terminus of the protein (^oe^*Tb*MfnL-ΔTM_::3Ty1_). **(H)** Western blotting of whole-cell extracts of *T. brucei* PCF overexpressing *Tb*MfnL-ΔTM_::3Ty1_ cells; (-Tet) non-induced and (+Tet) induced cells for 3 days. **(I)** Subcellular localization of ^oe^*Tb*MfnL-ΔTM_::3Ty1_ in PCF after 3 days induction, analyzed by standard immunofluorescence. n, nucleus; k, kinetoplast. The scale bar represents 5 µm.

To confirm the role and relevance of these domains on *Tb*MfnL localization and function, we overexpressed 3Ty1-tagged *Tb*MfnL-mutants without MTS (^oe^*Tb*MfnL-ΔMTS_::3Ty1_) or without TMs (^oe^*Tb*MfnL-ΔTM_::3Ty1_) in PCF trypanosomes (**Figure 7D and 7G**). Tetracycline-induced overexpression of both recombinant *Tb*MfnL was confirmed by western blot, 3 to 5 days after induction (**Figure 7E/7H, Figure S9**). In contrast to ^oe^*Tb*MfnL_::3Ty1_, ^oe^*Tb*MfnL-ΔMTS_::3Ty1_ did not colocalize with Tdh revealing that the N-terminal MTS is indeed required for proper mitochondrial targeting of *Tb*MfnL (**Figure 7F**). The MTS is therefore essential for the correct mitochondrial localization of *Tb*MfnL. In addition, overexpression of ^oe^*Tb*MfnL-ΔMTS_::3Ty1_ did not alter the mitochondrial structure, indicating that its proper, MTS-mediated, mitochondrial localization is essential for *Tb*MfnL-function (**Figure 7F/Anti-Tdh, Figure S9**). Overexpression of a *Tb*MfnL truncated version missing the C-proximal TMs (^oe^*Tb*MfnL-ΔTM_::3Ty1_, **Figure 7G-H, Figure S9**) did not affect its mitochondrial localization (**Figure 7I**), which confirms that the N-terminal MTS sequence was sufficient for mitochondrial targeting of *Tb*MfnL. However, in contrast to full length *Tb*MfnL (**Figures 4C and 6A/B**), *Tb*MfnL lacking TMs did not induce a modification of the mitochondrial shape (**Figure 7I, Figure S9**), demonstrating that the transmembrane domains are required for proper function. Altogether, our data showed that *Tb*MfnL is targeted to the mitochondrion via an N-terminal MTS and that C-proximal TMs are required for the cellular function of *Tb*MfnL.

Finally, we investigated the precise localization of *Tb*MfnL_::3Ty1_ and its variants (*Tb*MfnL-ΔTM_::3Ty1_ and *Tb*MfnL-ΔMTS_::3Ty1_) in the clone 2A11, where the endogenous *Tb*MfnL alleles were tagged with 3Ty1 (**Figure S5A-D**). This was achieved using a protease protection assay, a method previously used to determine the submitochondrial localization of various mitochondrial proteins in trypanosomes [44, 45]. Mitochondria are enriched by differential centrifugation, converted into mitoplasts by digitonin-mediated permeabilization of the outer membrane and finally incubated with proteinase K in the presence or absence of Triton X-100 to permeabilize the inner membrane.

Western blot analysis revealed that the cytosolic protein Enolase was almost entirely recovered in the supernatant, confirming that the generation of mitoplasts had effectively released cytosolic components (**Figure 8A, Figure S12**). In contrast, the *Tb*MfnL_::3Ty1_ protein and its variants exhibited fractionation patterns consistent with those of mitochondrial proteins, such as Atom40, Prohibitin, and Tdh, which serve as markers for the outer mitochondrial membrane (OMM), inner mitochondrial membrane (IMM, facing the intermembrane space), and mitochondrial matrix, respectively (**Figure 8A/C, Figure S12**). Proteinase K treatment of mitoplasts led to the near-complete digestion of outer membrane Atom40, intermembrane space Prohibitin (bound to the inner membrane), and *Tb*MfnL-ΔMTS_::3Ty1_, which is no longer targeted to the mitochondrion and thus unprotected. In contrast, the size and levels of *Tb*MfnL_::3Ty1_, *Tb*MfnL-ΔTM_::3Ty1_ and Tdh remained unaffected, indicating protease protection by the inner membrane. The intact state of Tdh further confirmed the integrity of the mitochondrial inner membrane and the reliability of the assay (**Figure 8A/C, Figure S12**). Digestion of the OMM-embedded β-barrel protein Atom40 and IMS Prohibitin, along with the preservation of *Tb*MfnL_::3Ty1_ and *Tb*MfnL-ΔTM_::3Ty1_, suggests that *Tb*MfnL faces the matrix and is protected by the inner membrane. The addition of Triton X-100, which disrupts membrane integrity, exposed matrix proteins to proteinase K, resulting in the degradation of *Tb*MfnL_::3Ty1_, *Tb*MfnL-ΔTM_::3Ty1_, and Tdh, thereby confirming their localization and orientation on the matrix side (**Figure 8A/C, Figure S12**). It should be noted that western blot revealed a size difference between the full-length *Tb*MfnL_::3Ty1_ and *Tb*MfnL-ΔTM_::3Ty1_. Specifically, we observed two bands: the upper band corresponding to the endogenously tagged protein and the lower band representing to the overexpressed *Tb*MfnL-ΔTM_::3Ty1_ protein.

**Figure 8.**
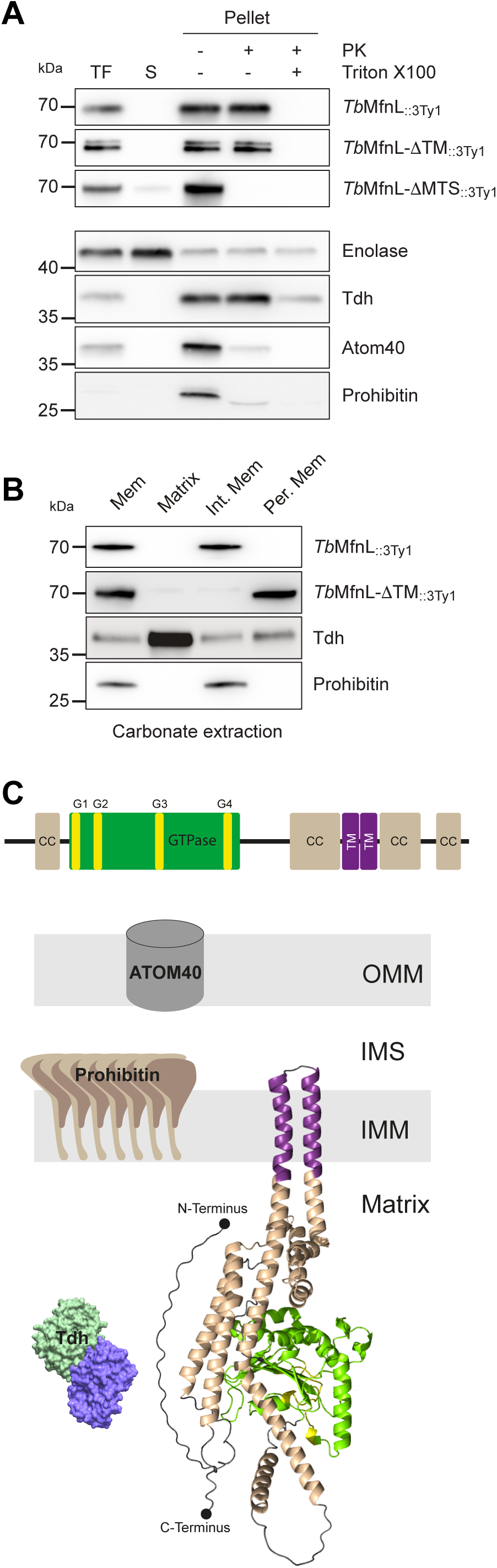
Membrane topology of *Tb*MfnL. **(A**) The Proteinase K protection assay involved incubating mitoplasts with Proteinase K and Triton X100, as denoted by the “+” symbol. TF represents the total fraction containing both cytoplasmic and membrane components; S denotes the supernatant, comprising the cytosolic contents; and Pellet represents the organelle fraction. **(B)** Carbonate extraction of mitochondria isolated from ^oe^*Tb*MfnL_::3Ty1_ and ^oe^*Tb*MfnL-ΔTM_::3Ty1_ cells. Mem, membrane fraction; Matrix, matrix fraction; Int. Mem, integral membrane fraction; Per. Mem, peripheral membrane fraction. **(C)** Predicted structure of *Tb*MfnL, from AlphaFold protein structure database. *Tb*MfnL was embedded in mitochondrial inner membrane (IMM). The color coding corresponds to the predicted secondary structure depicted on the top of the figure. The first 41 N-terminal amino acids corresponding to the MTS are not shown as they are cleaved. The N- and C-termini are marked by small black spheres. The AlphaFold model score of *Tb*MfnL is between confident to very high for most of the structured domains (https://alpha-fold.ebi.ac.uk/entry/Q57XN3, **Figure S13**).

To further investigate the localization of *Tb*MfnL, we separated the membrane and matrix fractions by performing repeated freeze-thaw lysis on mitoplast preparations. The membrane fraction was further divided into integral and peripheral fractions via carbonate extraction. *Tb*MfnL_::3Ty1_ and *Tb*MfnL-ΔTM_::3Ty1_ were analyzed alongside control proteins, Tdh (matrix-localized) and Prohibitin (inner membrane, IMS oriented). As expected, *Tb*MfnL_::3Ty1_ was exclusively retained in the membrane fraction, whereas *Tb*MfnL-ΔTM_::3Ty1_, lacking its transmembrane domains, was released from membrane (**Figure 8B/C, Figure S12**). These data are consistent with a recent analysis of the mitochondrial proteome, which proposed that the *Tb*MfnL protein is located in the inner mitochondrial membrane [46].

Interestingly, structural predictions of *Tb*MfnL using Alphafold, supported by a very high pLDDT score (**Figure S13**), suggest that the two previously identified transmembrane domains form a loop. This loop may act as an anchor within the inner membrane, thereby orientating the protein towards the matrix (**Figure 8C**), consistent with the findings from proteinase K assay and carbonate extraction.

## DISCUSSION

Membrane fusion and fission are often controlled by proteins of the dynamin superfamily (DSP) [47, 48]. These proteins consistently have a GTPase domain and contain domains required for membrane binding and oligomerization. In trypanosomes, only DSPs involved in fission have been characterized, and no fusion proteins similar to those of yeasts and mammals (Mfn and Opa1) have been identified so far. This may suggest that trypanosome mitochondria fuse via a machinery that differs from that characterized in yeasts and mammals. We have identified a novel DSP, named Mfn-Like (MfnL), which first representative characterized from the protozoan parasite *T. brucei* (*Tb*MfnL). We dit not establish a direct role in fusion, but demonstrated that it is able to increase mitochondrial interconnectivity in a GTPase-dependent manner, as previously reported for fusion factors Mfn/Opa1 [43, 49]. Interestingly, *Tb*MfnL domain organization and sub-mitochondrial localization differ significantly from that of Mfn and Opa1, suggesting that *Tb*MfnL and possibly other MfnL members mediate and/or enhance membrane branching and interconnection through a different molecular mechanism.

As observed for trypanosomes, many organisms including plants, lack genes encoding protein similar to known fusion factors. Interestingly, the search for homologous sequences to *Tb*MfnL allowed us to identify close DSP homologs in several eukaryotic phyla lacking Mfn and Opa1. These new DSPs are widely distributed in eukaryotes, which suggests that they represent an ancient protein family. Most of these phyla belong to the Diphoda, which together with the Opimoda are the two major supergroups of eukaryotes based on the most accepted position of the root of the tree of eukaryotes [50]. The only opimodan sequences found were the two amoeba cited above, most likely representing HGTs from bacteria, and the malawimonad *Gefionella okellyi* (**Figure 1C and Figure S3**). This malawimonad sequence may also reflect an HGT acquisition, in this case from a diphodan ancestor. The taxonomic distribution of this protein and its phylogeny support that it was ancestral at least in Diphoda. However, its presence also in bacteria, with any clear evidence of an HGT-mediated eukaryotic origin of the bacterial sequences, opens the possibility for an even more ancient origin before the diversification of Diphoda and Opimoda, which would imply a massive loss in opimodan taxa except in a few ones such as *Gefionella*. Future analysis of other deep-branching opimodan taxa will help to test this hypothesis. Homologs of *Tb*MfnL were also identified in several bacterial phyla, suggesting the existence in these organisms of a novel machinery involved in membrane shaping and/or remodeling that is different from the one found in yeasts and mammals. While studies of yeast and mammalian DSPs provide information about their functions and structures, little is known about bacterial DSPs, especially their cellular role. Indeed, some bacterial DSPs have been associated with a variety of processes involving membranes *in vivo*, such as a surveillance mechanism for membrane punctures caused by antibiotics and bacteriophages [51, 52], membrane vesicle formation [53] or cytokinesis by promoting membrane curvature at the septum [54, 55]. In addition, studies of the *Bacillus subtilis* DSP DynA have shown that this protein can promote membrane fusion *in vitro* [56]. The mechanism by which the fusion operates is not yet known, but bacterial DSPs are probably recruited to the sites where membrane fusion is needed [55, 57]. Interestingly, structural analysis of *Campylobacter jejuni* DLP1/DLP2 proteins allowed to propose a mechanism explaining how these proteins attach and bind distant and opposing membranes [58]. Unfortunately, the cellular function of *C. jejuni* DLP1/DLP2 is currently unknown. DSPs identified in this study have not yet been characterized and no information on their localization and function is available. However, among the identified organisms, we found bacteria of the phylum Planctomycetes, which are unusual organisms. Indeed, these bacteria contain a compartmentalized cytosol, separated by an intracytoplasmic membrane [59], in some cases surrounding the DNA [60] or forming an anammoxosome, a membrane-bound organelle responsible for the production of energy [61, 62]. It is then conceivable that the identified protein in Planctomycetes (*Pb*DBF, **Figure 1 and S2**) is involved in membrane structuring.

The role of *Tb*MfnL in mitochondrial shaping was observed by its overexpression in PCF trypanosomes, which induced a very strong increase in number of connections in the single branched mitochondrion. Moreover, the inactivation of its GTPase domain via mutation of the G1 motif also confirms its belonging to the large family of dynamins. Interestingly, a link between hyperconnection of a branching network and increased fusion processes has been reported in a highly different biological system, filamentous fungi, which grow as interconnected branching networks [63]. This raises the intriguing possibility that an excess of *Tb*MfnL in PCF trypanosomes enhances the interconnectivity of mitochondrial networks by inducing mitochondrial fusion.

The absence of fragmentation observed upon partial or total inactivation of *Tb*MfnL expression is in agreement with the absence or a very low level of constitutive mitochondrial fission. In contrast, in mammals and yeast, inhibition of fusion results in mitochondrial fragmentation by constitutive fission [16, 64]. Thus, our results are in agreement with previous studies suggesting that fission only occurs during cell division [19, 28]. The lack of observable mitochondrial alterations may also be attributed to the involvement of other proteins in mitochondrial structuring, as suggested by studies on the mitochondrial outer membrane proteome in *T. brucei* [65, 66] and the presence of mitochondrial complexes such as the recently characterized MICOS [44]. Additionally, subtle changes in mitochondrial structure may exist but could have gone undetected using our current approaches. It is worth noting that *Tb*MfnL seems to participate in a unique membrane-structuring mechanism, and its inactivation may not produce effects similar to those observed for Mfn or Opa1.

Another intriguing point was the discrepancy with the results reported by Vanwalleghem et al. [30]. Indeed, they describe fenestration of the mitochondrion in BSF cells with down-regulated *Tb*MfnL, whereas in our studies, involving inactivation by CRISPR/Cas9 or down-regulation by RNAi in BSF cells, we did not observe any alterations in mitochondrial structure. The only modification (hyper-reticulation) of the mitochondrial structure was observed exclusively during the overexpression of *Tb*MfnL and only in PCF forms. This discrepancy may arise from differences in cell lines, culturing conditions, sample preparation methods, or mitochondrial staining protocols. We used rhodamine staining on live cells and immunofluorescence labeling to visualize the mitochondrial network, whereas Vanwalleghem et al. [30] employed FIB-SEM tomography. It is also possible that *Tb*MfnL knockdown affects mitochondrial import, since disruptions in mitochondrial import have been reported to influence mitochondrial structure [65].

Using several microscopy approaches, we have demonstrated that *Tb*MfnL is localized within the mitochondrial membranes, with two domains involved in its accurate mitochondrial addressing, a canonical N-terminal MTS and two adjacent C-terminal transmembrane domains. Expression of MTS-less *Tb*MfnL or MTS-GFP recombinant proteins in PCF trypanosomes confirmed the role of this 41 aa-long MTS for mitochondrial import. Proteins with an N-terminal MTS are typically imported into the mitochondrial matrix, with a few exceptions, such as Opa1 and Apoptosis-inducing factor (AIF) face the intermembrane space [67]. Indeed, Opa1 and AIF localize mainly to the IMS due to their TM domains located in close proximity to the MTS (**Figure 1A**), interrupting transport across the inner membrane. The C-proximal localization of the transmembrane domains of *Tb*MfnL is not compatible with such import interruption, but more presumably suggested that its C-terminus is also exposed to the matrix, as predicted by Alphafold (**Figure 8C**). This model is consistent with the proteinase K resistance of *Tb*MfnL in mitoplasts. This orientation is also probably similar to that of the prokaryotic MfnLs identified in this study, which could also be anchored in the plasma membrane and orientated towards the cytosol. Interestingly, recent publications describe mitochondrial remodeling originating from the matrix side of the mitochondria. Sheikh *et al.* [68] demonstrated that expressing a potential DSP from a giant virus in *T. brucei* PCF significantly affects mitochondrial morphology within the matrix, with a notable association with the inner membrane. Additionally, Kumar *et al.* [69, 70] showed that a dynamin superfamily-like pseudoenzyme, a distant relative of the dynamin superfamily, stabilizes cristae architecture through interactions on the matrix side of the mitochondrial inner membrane. This contrasts with the mode of action of human or yeast mitofusins, which are involved in the remodeling of mitochondrial outer and inner membranes, but never via the matrix side.

In conclusion, the data presented here uncover a new family of DSPs and suggest the existence of a novel membrane-structuring mechanism found in both eukaryotes and prokaryotes, but distinct from those involved in mitochondrial membrane fusion in mammals and fungi.

## ACKNOWLEDGMENTS AND FUNDING SOURCES

We thank Hassan Hashimi (University of South Bohemia) for providing us the anti-Prohibitin antibody, André Schneider (University of Bern) for providing us the anti-Atom40 antibody, Keith Gull (University of Manchester) for providing us the anti-Ty1 antibody and Derrick Robinson, Mélanie Bonhivers, Elina Casas and Nicolas Landrein (University of Bordeaux) for the pPOTv7 expression vector and extremely valuable help in the expansion microscopy experiment. Cell sorter analyses were performed at the TBMCore facility (FACSility) on BD FACSAria™ III Sorter and we thank Atika Zouine for data acquisition and interpretation. We also thanks Bordeaux Imaging Center (BIC), which is a member of the FranceBioImaging national infrastructure (ANR10-INBS-04), for helping us to design the ImageJ macro and microscopy acquisitions. The Bringaud team is supported by the Centre National de la Recherche Scientifique (CNRS, https://www.cnrs.fr/), the Université de Bordeaux (https://www.u-bordeaux.fr/) and the Agence Nationale de la Recherche (ANR, https://anr.fr/) through the ParaFrap “Laboratoire d’Excellence” (LabEx, https://www.enseignementsup-recherche.gouv.fr/cid51355/laboratoires-d-excellence.html) (ANR-11-LABX-0024). This work was supported by the “Fondation pour la Recherche Médicale” (FRM, https://www.frm.org/) (“Equipe FRM”, grant n°EQU201903007845) and the ANR grant ADIPOTRYP (ANR19-CE15-0004-01) to FB and the ERC Advanced grant 787904 to D.M. Portions of this manuscript were based on Dr. Morel’s thesis defended at the University of Bordeaux.

## AUTHOR CONTRIBUTIONS

C.A.M. conceptualized the study, performed experiments, contributed to analysis and writing. C.A. performed experiments and contributed to analysis. D.M. performed all the phylogenic analysis. C.B., B.S. and E.G. performed the EM and SBF-SEM experiments and contributed to their analysis. S.D-C., M.R., F.B. and E.T. conceptualized the study, contributed to analysis and wrote the manuscript, with also an input from all the authors.

## MATERIALS AND METHODS

### Trypanosomes and cell cultures

The PCF of *T. brucei* EATRO1125.T7T (TetR-HYG-T7RNAPOL-NEO, where TetR stands for tetracycline resistance, HYG for hygromycin, T7RNAPOL for RNA polymerase T7, and NEO for neomycin) was cultured at 27°C with 5% CO_2_ in SDM79 medium containing 10% (vol/vol) heat-inactivated fetal calf serum, 5 µg/ml hemin, 25 µg/ml hygromycin and 10 µg/mL neomycin. PCF cells were also cultured in SDM80 medium containing either glucose (10 mM) as the sole carbon source or proline (5 mM) supplemented with N-acetyl-glucosamine (50 mM) to inhibit the glucose transporter. The bloodstream form of *T. brucei* 427 90-13 (TetR-HYG-T7RNAPOLNEO) was cultured at 37°C with 5% CO_2_ in Iscove’s modified Dulbecco’s medium (IMDM) supplemented with 10% (vol/vol) heat-inactivated fetal calf serum, 0.2 mM Δ-mercaptoethanol, 36 mM NaHCO_3_, 1 mM hypoxanthine, 0.16 mM thymidine, 1 mM sodium pyruvate, 0.05 mM bathocuproine, 1.5 mM L-cysteine [74], 5 µg/ml hygromycin and 2.5 µg/mL neomycin. Overexpression cell lines were induced with tetracycline (10 µg/mL for BSF and 1 µg/mL for PCF). Growth was monitored by daily cell counting with the cytometer Guava® Muse® and Guava® easyCyte™.

### Endogenous tagging and inactivation of *Tb*MfnL by CRISPR-Cas9

Endogenous tagging and inactivation of *Tb*MfnL were achieved by CRISPR Cas9 technology [36]. Briefly, inactivation of the *Tb*MfnL was achieved by inserting the resistant marker puromycin (Pac) or a small sequence encoding for a *Bam*HI restriction site plus 6 successive stop codons, flanked by 50 bp homologous to the 5’ and 3’ *Tb*MfnL sequences from the Cas9 cutting site. Similar approaches have been performed for endogenous tagging with the insertion of a sequence encoding 3xTy1 or 3xHA. The EATRO1125.T7T PCF and the 427 90.13 BSF (1×10^6^ cells) were respectively transfected, using Amaxa nucleofectorII, with 1 µg of purified cassette (puromycin resistance marker, Stop*Bam*H1Stop, 3xTy1 or 3xHA), 30 µg Cas9 protein from IDT and pre-annealing TracrRNA (0.4 µmol) and gRNA (0.4 µmol). Cells were transfected using program X-001 and selected or not with puromycin (SDM79, 1 µg/mL, or IMDM 0.1 µg)/mL). Cells were cloned by using a cell sorter (TBM Core facility), and selection of double inactivated *Tb*MfnL gene (*Tb*MfnL^−/-^) or endogenously tagged clones was done by DNA extraction, with NucleoSpin Blood (Macherey-Nagel) and PCR amplification, see supplemental **Table S2**. Guide RNAs were designed using EuPaGDT, from http://tritrypdb.org. Primers and guide RNAs used were synthetized by Integrated DNA Technologies (IDT) and are listed in supplemental **Table S2**.

### Immunofluorescence

Cells were washed twice with PBS, then fixed with 2% paraformaldehyde (PFA) for 10 min at room temperature and 0.1mM glycine was added 10 min to stop the reaction. The cells were spread on slides and permeabilized with 0.05% triton X-100. After incubation in PBS containing 4% bovine serum albumin (BSA) 20 min, cells were incubated for 1 hour with primary antibodies diluted in PBS-BSA 4%, washed 4 times with PBS and incubated for 45 min with secondary antibodies diluted in PBS-BSA 4% followed by three washes. Then, kinetoplasts and nuclei were labelled with DAPI (10 µg/mL) for 5 min. Slides were washed 3 times with PBS and mounted with SlowFade Gold (Molecular probes). Images were acquired with MetaMorph software on Zeiss Imager Z1 or an Axioplan 2 microscope and processed with ImageJ.

### Mitochondria staining in live cells

Rhodamine-123 (30 µg/mL) was added on cell culture (5×10^6^ - 1×10^7^ cells per mL) for 15 min at room temperature, then cells were washed twice with PBS and spread on slides. As an alternative to labeling mitochondria in living cells with Rhodamine-123, Mitotracker (MitoTracker Green FM) was used. Cells were washed once with PBS, incubated for 10 minutes with 10 µM MitoTracker Green FM, washed again with PBS, and then spread onto slides. Images were acquired with MetaMorph software on an Axioplan 2 microscope and processed with ImageJ.

### Ultrastructure Expansion Microscopy (UExM)

The protocol of UExM was realized exactly as described by Casas et *al.* 2022 (dx.doi.org/10.17504/protocols.io.bvwqn7dw). An expansion factor was determined using the ratio between the size of the coverslip (12 mm) and the size of the gels after the first expansion. Images were acquired on a confocal Leica SP5-MP (Bordeaux Imaging Center) using a 63X oil objective and processed with ImageJ.

### Quantification of Mitochondrial junctions - ImageJ macro

To quantify the number of mitochondrial junctions, an ImageJ macro was developed with the help of the Bordeaux Imaging Center (BIC). The script is presented in supplemental (**Data S1**).

### Endogenous tagging and overexpression of *Tb*MfnL

For endogenous gene tagging, primers were designed as described in [75] and PCR was performed using pPOTv7 vector as template. The pPOT used here was pPOTv7 for C-terminus 10xHA tagging (blasticidin resistance). For overexpression, *MfnL* gene (Tb427.7.2410) was inserted in both pLew100 (phleomycine resistance) [76] and pHD1336 (blasticidin resistance) [77] expression vectors. Fragments were amplified and cloned into *Hin*dIII and *Xb*aI restrictions sites of the pLew100 and into *Hin*dIII and *Bam*HI restrictions sites of the pHD1336 expression vectors. In addition, a 3xTy1 tag was added at the C-terminus of the protein in the pLew100-*Tb*MfnL, using the *Xba*I and *Bam*HI restrictions sites. The truncated version of *Tb*MfnL without the first 41 amino acids (*Tb*MfnL-DMTS) and without the two trans-membranes domain (*Tb*MfnL-DTM) were amplified and cloned in the pLew100 with a 3xTy1 tag as previously described. Catalytic lysine (K141) was replaced by an alanine, using PCR approach [78] and PCR product cloned in the pLew100 with a 3xTy1 tag (*Tb*MfnL-K141A). EATRO1125.T7T PCF was transfected using Amaxa Nucleofector®II, program X-001 and selected in SDM79 containing 25 µg/ml hygromycin 10 µg/mL neomycin and 5 µg/mL phleomycin. 427 90-13 BSF was transfected using the same conditions and selected in IMDM containing 5 µg/ml hygromycin, 2.5 µg/mL neomycin and 5 µg/mL blasticidin. Primers used for the constructions are presented in supplemental **Table S2**.

### Down-regulation of *Tb*MfnL gene expression

Down-regulation of *Tb*MfnL expression by RNAi in PCF was achieved by expression of stemloop “sense/antisense” RNA molecules targeting a 400-bp fragment of the *Tb*MfnL gene introduced into the pLew100 tetracycline-inducible expression vector. A PCR-amplified 450-bp fragment, containing the antisense *Tb*MfnL sequence was inserted between *Xho*I and *Bam*HI restriction sites of the pLew100 plasmid. Then, the separate 400-bp PCR-amplified fragment containing the sense *Tb*MfnL sequence was inserted upstream of the antisense sequence, using *Hin*dIII and *Xho*I restriction sites. The resulting plasmid, pLew100-*Tb*MfnL-SAS, contains a sense and antisense version of the *Tb*MfnL fragment separated by a 50-bp fragment. The EATRO1125 PCF was transfected with the pLew100-*Tb*MfnL-SAS and cells were selected in SDM79 medium containing 25 µg/ml hygromycin, 10 µg/mL neomycin and 5 µg/mL phleomycin. Expression of the RNAi was induced by tetracycline (1 µg/mL). Down-regulation of *Tb*MfnL expression by RNAi in BSF was achieved exactly as described [30] with a 280-bp fragment derived from the *Tb*MfnL open reading frame inserted into the p2T7-177 plasmid [79]. Linearized plasmid was transfected in *T. brucei* 427 90-13 (TetR-HYG-T7RNAPOL-NEO) BSF cells. Expression of the RNAi was induced by tetracycline (10 µg/mL). Primers used for the constructions are presented in supplemental **Table S2**.

### Serial Block-Face Scanning Electron Microscopy (SBF-SEM)

The entire procedure is detailed and has been carried out as described in the article: Advancing yeast cell analysis: A cryomethod for serial block-face scanning electron microscopy imaging in mitochondrial morphology studies [80].

### Immunoelectron microscopy

Harvested cells were placed on the surface of formvar-coated copper grids (400 mesh). Each loop was quickly submersed in liquid propane (−180°C) and transferred into a precooled solution of 0.1% uranyl acetate in dry acetone for 3 days at -82°C. Samples were rinsed with acetone at -20°C, and embedded progressively at -20°C in LR Gold resin (EMS, USA). Resin polymerization was carried out at -20°C for 7 days under UV illumination. Ultrathin LR Gold sections were collected on nickel grids coated with formvar. Sections were first incubated for 15 min with 500mM NH_4_Cl in Tris-buffered saline (TBS) pH7.8, blocked 2×10 min with 2% BSA in TBS pH 7.8. The grids were incubated 1 hour at room temperature with anti Ty1 antibody diluted to 1:200 in TBS containing 2% BSA rinsed with TBS containing 2% BSA and with TBS containing 1% BSA. The samples were then incubated for 1 hour at room temperature with anti-mouse IgG diluted to 1:20 conjugated to 10 nm gold particles (BioCell). The sections were rinsed with TBS containing 1% BSA, and fix with 1% glutaraldehyde in TBS. After rinsing with TBS, grids were contrasted through a 5 min incubation with 2% uranyl acetate in water, followed by 1 min incubation with 1% lead citrate. Observations were performed on a HITACHI H7650 transmission electron microscope operated at 80 KV with an Orius 1000-11 MPx camera (Gatan, Abingdon, UK).

### Transmission electron microscopy (TEM)

The entire procedure is thoroughly detailed and has been carried out precisely as described in the article: Plunge Freezing: A Tool for the Ultrastructural and Immunolocalization Studies of Suspension Cells in Transmission Electron Microscopy [81].

### Western blot analyses

Total protein extracts (5×10^6^ cells) were separated by SDS-PAGE (10%) and immunoblotted on TransBlot Turbo Midi-size PVDF Membranes (Bio-Rad). Immunodetection was performed using the primary antibodies, diluted in PBS-Tween Milk (0.05% Tween20, 5% skimmed milk powder), summarized in supplemental **Table S3**. Revelation was performed using a second antibody coupled to the horseradish peroxidase and the Clarity western enhanced-chemiluminescence (ECL) substrate as describes by the manufacturer (Bio-Rad). Images were acquired and analyzed with the ImageQuant Las 4000 luminescent image analyzer.

### Mitochondria-enriched preparation, proteinase K protection assays and carbonate extraction

This procedure has been well established and allows the determination of the submitochondrial localization of trypanosome proteins [44, 45, 82]. 10^8^ cells were lysed in SoTE buffer (20 mM Tris-HCl, pH 7.5, 0.6 mL sorbitol, and 2 mM ethylenediaminetetraacetic acid), supplemented with 0.02% (w/v) digitonin (D141-100MG, Sigma-Aldrich) for 5 min at 4°C. A portion of the resulting suspension was collected as the total fraction (TF), while the remaining suspension underwent centrifugation at 6,000 x g for 3 minutes at 4°C to isolate a mitochondria-enriched pellet. This pellet was then resuspended in SoTE supplemented with 0.2% (w/v) digitonin and incubated on ice for 15 minutes. The resuspended mixture was divided into three tubes and subjected to centrifugation at 6,000 × g for 3 minutes at 4°C, resulting in the separation of mitoplasts and soluble protein in the supernatant (S). Each mitoplast pellet was resuspended in SoTE and subjected to the proteinase K protection assay. Proteinase K (50 µg) was added to two tubes, one supplemented with 1% Triton-X100 (v/v), while the third tube remained untreated as a control. All tubes were then incubated on ice for 30 minutes, followed by addition of 5 mM phenylmethylsulfonyl fluoride to stop the reaction. Proteins from all fractions were precipitated with trichloroacetic acid before undergoing resolution by SDS-PAGE.

For the extraction of integral membrane proteins using the carbonate method, the procedure begins with a mitochondria-enriched pellet. The pellet is resuspended in 500 µL of 10 mM MgCl₂ and subjected to a freeze-thaw cycle repeated ten times, involving freezing in liquid nitrogen and thawing at room temperature. Subsequently, the suspension is divided into two fractions: 50 µL and 450 µL. The 450 µL fraction is centrifuged at 10,000 × g for 5 minutes at 4 °C, with the supernatant designated as the matrix fraction. The 50 µL fraction is centrifuged under the same conditions, and the resulting pellet is resuspended in 50 µL of 10 mM MgCl₂, representing the membrane fraction. The remaining pellet from the 450 µL sample is further processed by resuspending it in 400 µL of 0.1 M Na₂CO₃ (pH 11). This mixture is incubated on ice for 30 minutes, sonicated four times, and centrifuged at 20,000 × g for 20 minutes at 4 °C. The supernatant is collected as the peripheral membrane protein fraction. The pellet is washed by resuspension in 400 µL of 0.1 M Na₂CO₃ and subjected to another centrifugation under the same conditions. This final pellet constitutes the integral membrane protein fraction. All protein fractions are precipitated using trichloroacetic acid, resuspended in 50 to 100 µL of 2% SDS, and analyzed by SDS–PAGE.

### Sequence database searches and phylogenetic analysis

Two databases were used to identify sequences homologous to the *T. brucei Tb*MfnL sequence (Tb927.7.2410). The first search was conducted using Blastp [83] (protein-protein BLAST, https://blast.ncbi.nlm.nih.gov/Blast.cgi) on the NCBI nr protein database, with *Tb*MfnL as the query and kinetoplastids excluded from the search. The parameters included a maximum of 5000 target sequences and the Blosum62 matrix. For the EuKProt v3 database [84] search (https://evocellbio.com/eukprot/), the Blastp server was used with the entire database (993 species) and *Tb*MfnL as the query, without applying advanced parameters. The selection of sequences was then performed manually, retaining only those sequences that exhibited homology across the entire *Tb*MfnL protein. Sequences corresponding solely to the highly conserved GTPase domain were excluded. A total of 315 sequences were selected for phylogenetic analysis. An additional 23 sequences were included from well-characterized dynamins involved in fission (Dnm1), fusion (Opa1, Mfn1/2, Fzo1), BDLP (the cyanobacterial Mfn homolog), and dynamins associated with clathrin-mediated endocytosis (Dyn1/2) to be used as an outgroup.

The sequences were aligned with MAFFT L-INS-i v7.450 [85] and trimmed with BMGE v1.12 (-m BLOSUM30 -b 3 -g 0.2 -h 0.5) [86]. A maximum likelihood phylogenetic tree was then reconstructed with IQ-TREE v.2.0.3 [87] using its model finder (-m TEST) option, including complex mixture models, to select the best fit substitution model, as well as 1000 ultrafast bootstrap replicates to estimate branch statistical support.

### Structural prediction software and sequence analysis

Structure predictions were performed using PSIPRED (“http://bioinf.cs.ucl.ac.uk/psipred/”) [72] and prediction of mitochondrial targeting sequences by various algorithms (MitoFates, PSORT II and TargetP-2.0) [88].

### Statistical analysis

Experiments were performed at least in triplicates. Statistical analyses were performed using Prism (GraphPad) software. The results are presented as mean ± S.D. Where indicated the results were subjected to two-sided Student’s t-test to determine statistical differences against the indicated group (Confidence interval 95% - P-value style: 0.1234 (ns); 0.0332 (*); 0.0021 (**); 0.0002 (***); <0.0001 (****)).

## SUPPLEMENTAL INFORMATION

### Data S1 - Code ImageJ: Mitochondrial junctions (MJ)

~~~
run(“Duplicate…”, “title=ORI duplicate channels=2”);
run(“Duplicate…”, “title=Estfond”);
run(“Gaussian Blur…”, “sigma=10”);
imageCalculator(“Divide create 32-bit”, “ORI”,“Estfond”);
selectWindow(“Result of ORI”);
rename(“OriCorr”); selectWindow(“Estfond”);
setAutoThreshold(“Li dark”);
run(“Create Selection”);
selectWindow(“OriCorr”);
run(“Restore Selection”);
run(“Enlarge…”, “enlarge=10 pixel”);
setAutoThreshold(“Li dark”);
run(“Analyze Particles…”, “size=15-500 show=Masks”);
rename(“Mask”);
run(“Options…”, “iterations=1 count=4 pad do=Open”);
run(“Duplicate…”, “title=Skeleton”);
setOption(“BlackBackground”, false);
run(“Skeletonize”);
run(“Duplicate…”, “title=Ends”);
run(“Options…”, “iterations=1 count=7 pad do=Erode”);
imageCalculator(“Subtract create”, “Skeleton”,“Ends”);
selectWindow(“Result of Skeleton”);
rename(“Endpoints”);
selectWindow(“Skeleton”);
run(“Duplicate…”, “title=Jonctions”);
run(“Options…”, “iterations=1 count=6 pad do=Erode”);
run(“Analyze Particles…”, “size=0-5000 summarize”);
imageCalculator(“Add create”, “Endpoints”,“Jonctions”);
selectWindow(“Result of Endpoints”);
rename(“Combo”);
imageCalculator(“Subtract create”, “Skeleton”,“Combo”);
selectWindow(“Result of Skeleton”);
rename(“Segments”);
selectWindow(“Combo”);
close();
selectWindow(“ORI”);
setOption(“ScaleConversions”, true);
run(“8-bit”);
run(“Merge Channels…”, “c1=Jonctions c2=Endpoints c3=Segments c4=ORI create ignore”); close(“\\Others”);
~~~

**Table S1.**
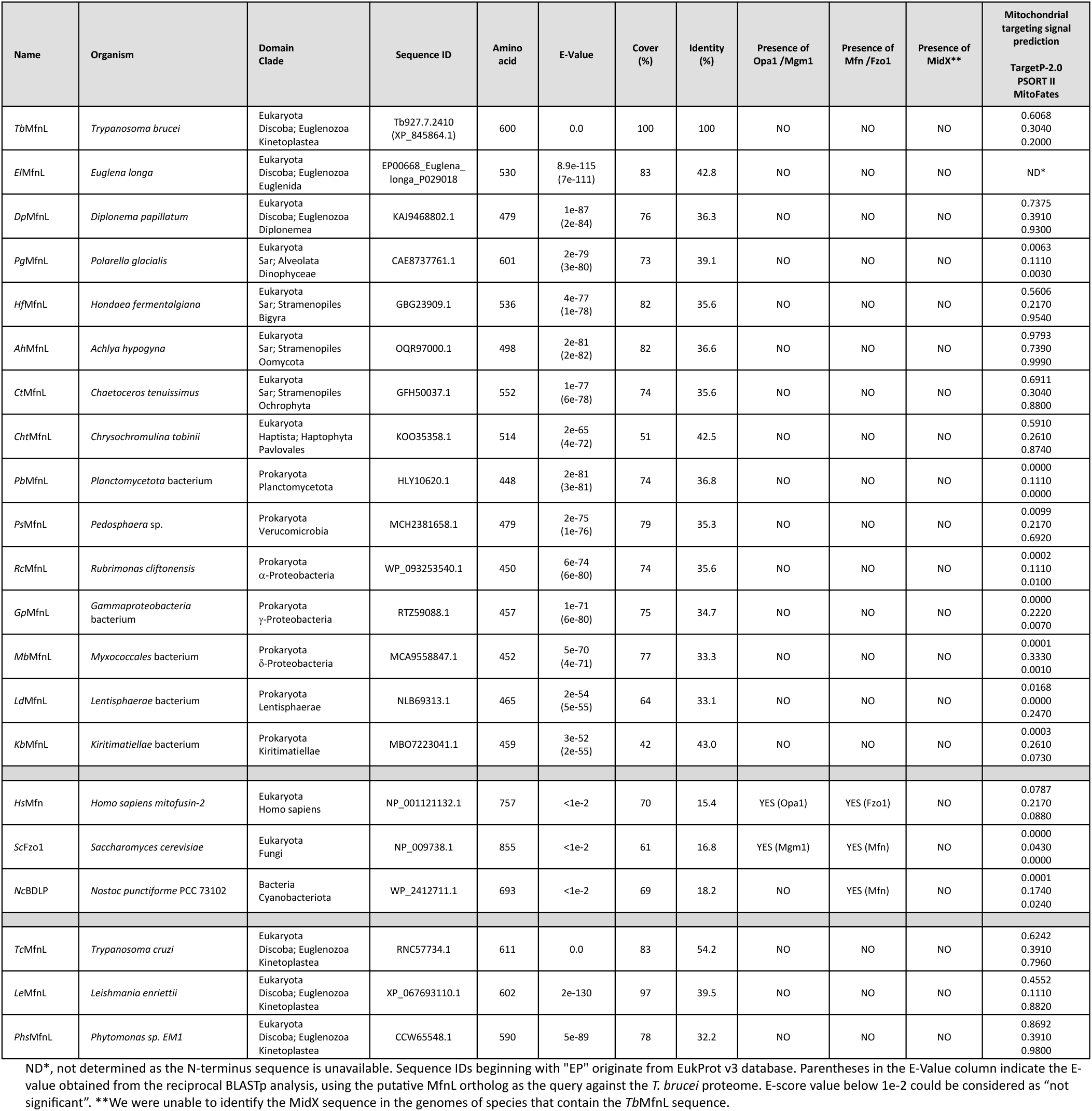
Results from BLASTp and EukProt v3 (query: *Tb*MfnL) of several identified organisms.

**Table S2.**
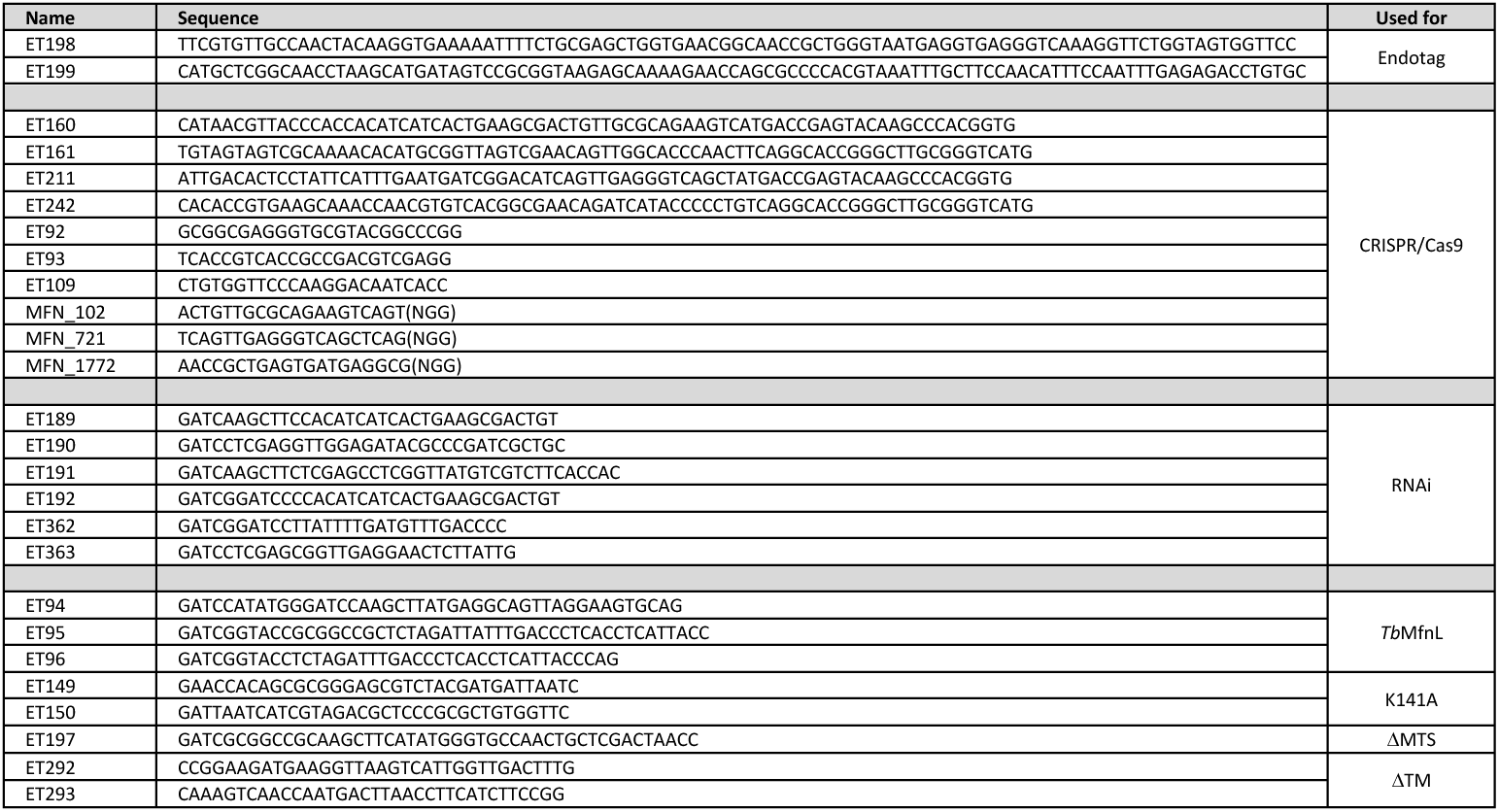
Primers used in this study.

**Table S3.** Antibodies used in this study.

| Name | Species | Dilution | Source/Ref |
| --- | --- | --- | --- |
| Anti-Ty1 | Mouse | WB 1:800 | iMET lab |
|  |  | IF 1:500 |  |
|  |  | UExM 1:250 |  |
| Anti-HA | Mouse | WB 1:1000 | Biolegend |
|  |  | IF 1:1000 |  |
|  |  | UExM 1:500 |  |
| Anti-HA | Rat | IF 1:1000 | Roche |
| Anti-Tdh | Rabbit | WB 1:500 | iMET lab |
|  |  | IF 1:200 |  |
|  |  | UExM 1:100 |  |
| Anti-Enolase | Rabbit | WB 1:100000 | Gift from P. Michels, Edinburgh, UK |
| Atom40 | Rabbit | WB 1:10000 | Gift from A. Schneider, Bern, Switzerland |
| Anti-Frdg | Rabbit | WB 1:50 | iMET lab |
| Anti-Hsp60 H7 | Mouse | IF 1:500 | iMET lab |
| Anti-PPDK | Rabbit | WB 1:1000 | iMET lab |
| Anti-GFP | Rabbit | WB 1:1000 | Abcam |
| Anti-Prohibitin | Rabbit | WB 1:1000 | Gift from H. Hashimi, Czech Republic |
| Anti-Mouse HRP | Goat | WB 1:5000 | Bio-Rad |
| Anti-Rabbit HRP | Goat | WB 1:10000 | Bio-Rad |
| Anti-Mouse Alexa Fluor 594 | Donkey | IF 1:400 | ThermoFisher |
|  |  | UExM 1:250 |  |
| Anti-mouse Alexa Fluor 488 | Donkey | IF 1:400 | ThermoFisher |
|  |  | UExM 1:250 |  |
| Anti-Rabbit Alexa Fluor 594 | Donkey | IF 1:400 | ThermoFisher |
|  |  | UExM 1:250 |  |
| Anti-Rabbit Alexa Fluor 488 | Donkey | IF 1:400 | ThermoFisher |
|  |  | UExM 1:250 |  |
| Anti-Rat Alexa Fluor 594 | Donkey | IF 1:400 | ThermoFisher |

**Figure S1.**
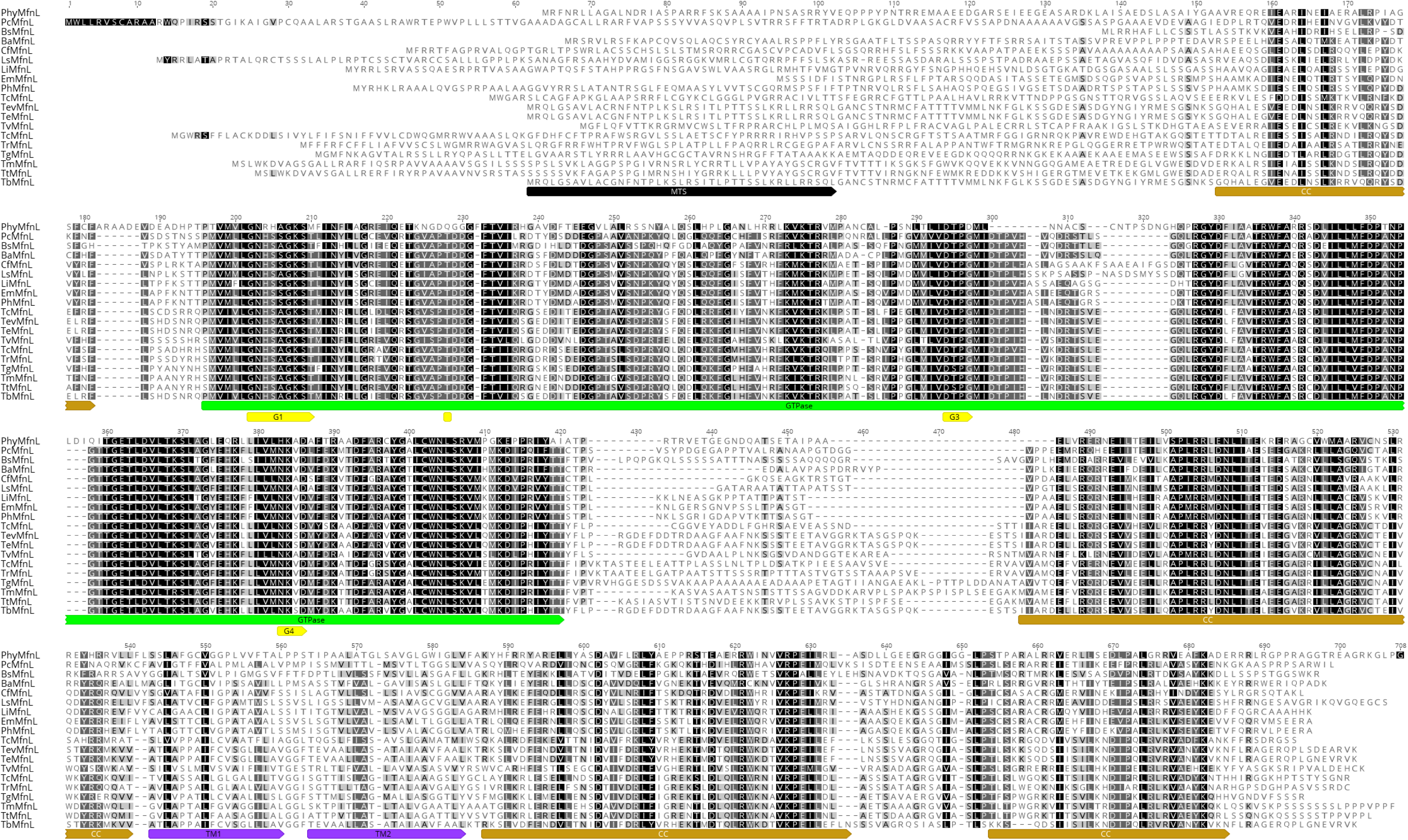
Sequence alignement between *Tb*MfnL and various kinetoplastids MfnL. MTS, predicted mitochondrial targeting sequence are indicated by grey boxes; GTPase domain is indicated by a green box, and the four GTP-binding motifs are indicated by yellow boxes and numbered; The coil coil domain are indicated by light brown boxes. TM, transmembrane span are indicated by purple boxes. Identities and similarities are represented by white characters on a black background and black characters on a grey background respectively. *Tb*MfnL, *Trypanosoma brucei* DRP (ID: Tb927.7.2410); *Phy*MfnL, *Phytomonas* sp. EM1(ID: CCW65548); *Pc*MfnL, *Paratrypanosoma confusum* (ID: PCON_0070060); *Bs*MfnL, *Bodo saltans* (ID: BSAL_56690); *Ba*MfnL, *Blechomonas ayalai* (ID: Baya_002_1420); *Cf*MfnL, *Crithidia fasciculata* (ID: CFAX1_240011500); *Ls*MfnL, *Leptomonas seymouri* (ID: Lsey_0356_0020); *Li*MfnL, *Leishmania infantum* (ID: LINF_220010300); *Em*MfnL, *Endotrypanum monterogeii* (ID: EMOLV88_220009500); *Ph*MfnL, *Porcisia hertigi* (ID: JKF63_04376); *Tc*MfnL, *Trypanosoma cruzi* (ID: TcCLB.508321.60); *Tev*MfnL, *Trypanosoma evansi* (ID: TevSTIB805.7.2480); *Te*MfnL, *Trypanosoma equiperdum* (ID: TEOVI_000623300); *Tv*MfnL, *Trypanosoma vivax* (ID: TvY486_0702250); *Tc*MfnL, *Trypanosoma congolense* (ID: TcIL3000_7_1710); *Tr*MfnL, *Trypanosoma rangeli* (ID: TRSC58_06192); *Tg*MfnL, *Trypanosoma grayi* (ID: DQ04_08691020); *Tm*MfnL, *Trypanosoma melophagium* (ID: LSM04_004715); *Tt*MfnL, *Trypanosoma theileri* (ID: TM35_000191450).

**Figure S2.**
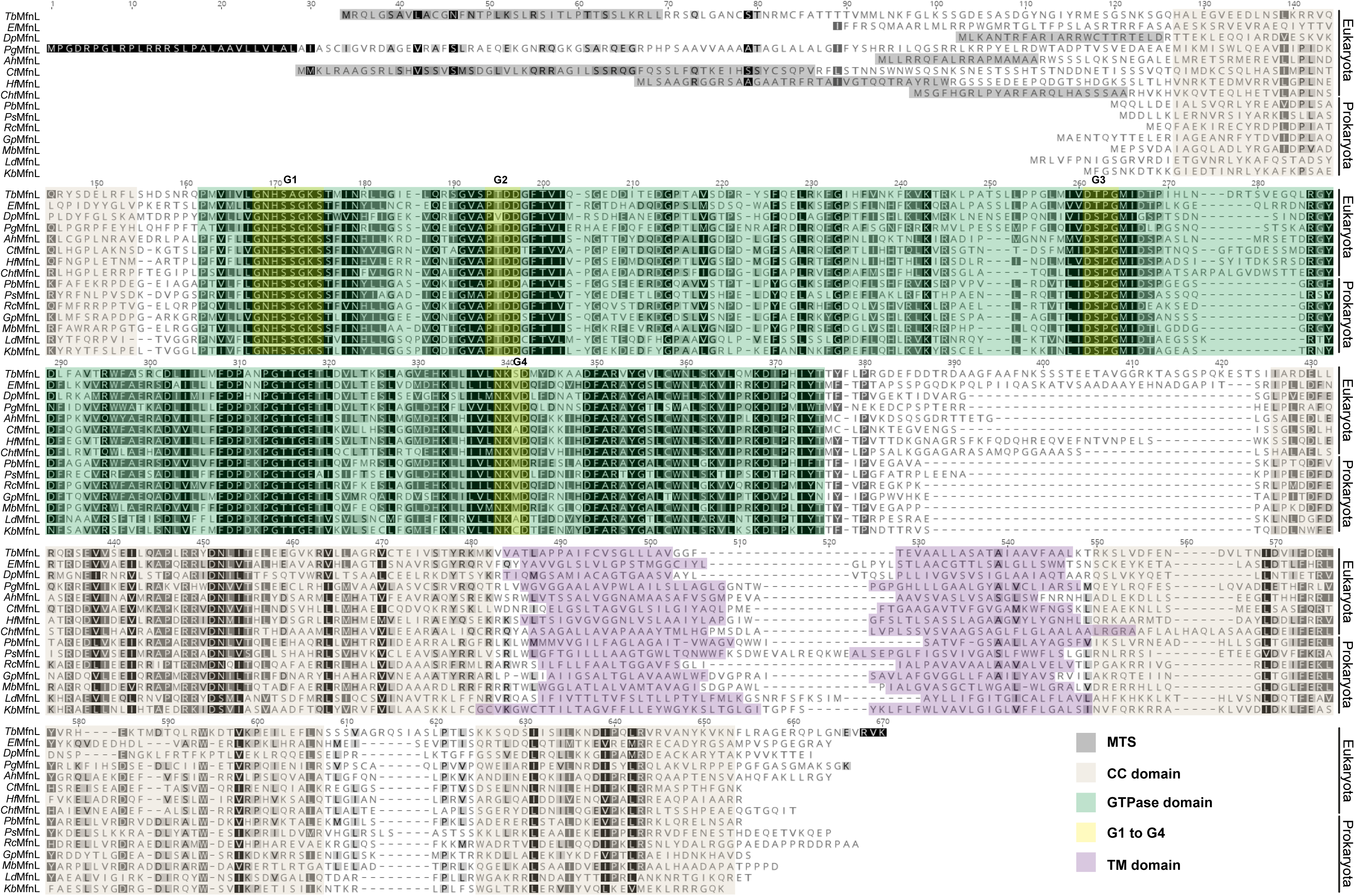
Sequence and structure alignement between *Tb*MfnL and homologous proteins identified in various eukaryotes and prokaryotes with BLASTp. MTS, predicted mitochondrial targeting sequence are highlighted in grey; GTPase domain is highlighted in green, and the four GTP-binding motifs are highlighted in yellow and numbered; The coil coil domain are highlighted in light brown. TM, transmembrane span are highlighted in purple. Identities and similarities are represented by white characters on a black background and black characters on a grey background respectively. Structure predictions were performed using PSIPRED and mitochondrial targeting sequences were predicted by various algorithms (TargetP2.0, PSORTII, MitoFates, MitoprotII and iPSORT). **Eukayota:** *Tb*MfnL, *Trypanosoma brucei* DRP (ID: Tb927.7.2410); *El*MfnL, *Euglena longa* (ID: EP00668_Euglena_longa_P029018); DpMfnL, *Diplonema papillatum* (ID: KAJ9468802.1)*; Pg*MfnL, *Polarella glacialis* (ID: CAE8737761.1); *Ah*MfnL, *Achlya hypogyna* (ID: OQR970001); *Ct*MfnL, *Chaetoceros tenuissimus* (ID: GFH50037.1); *Hf*MfnL, *Hondaea fermentalgiana* (ID: GBG23909.1); *Cht*MfnL, *Chrysochromulina tobinii* (ID: KOO35358.1). **Prokaryota:** *Pb*MfnL, *Planctomycetaceae* bacterium (ID: MBV8881356.1); *Ps*MfnL, *Pedosphaera* sp. (ID: MCH2381658.1); *Rc*MfnL, *Rubrimonas cliftonensis* (ID: WP_093253540.1); *Gp*MfnL, *Gammaproteobacteria* bacterium (ID: RTZ59088.1); *Mb*MfnL, *Myxococcota* bacterium (ID: MBU0550280.1); *Ld*MfnL, *Lentisphaerae* bacterium (ID: NLB69313.1); *Kb*MfnL, *Kiritimatiellae* bacterium (ID: MBO7223041.1). The accession number for *Euglena longa* was obtained from Eukprot V3 database. The alignment was performed using Geneious software with the BLOSUM62 matrix.

**Figure S3.**
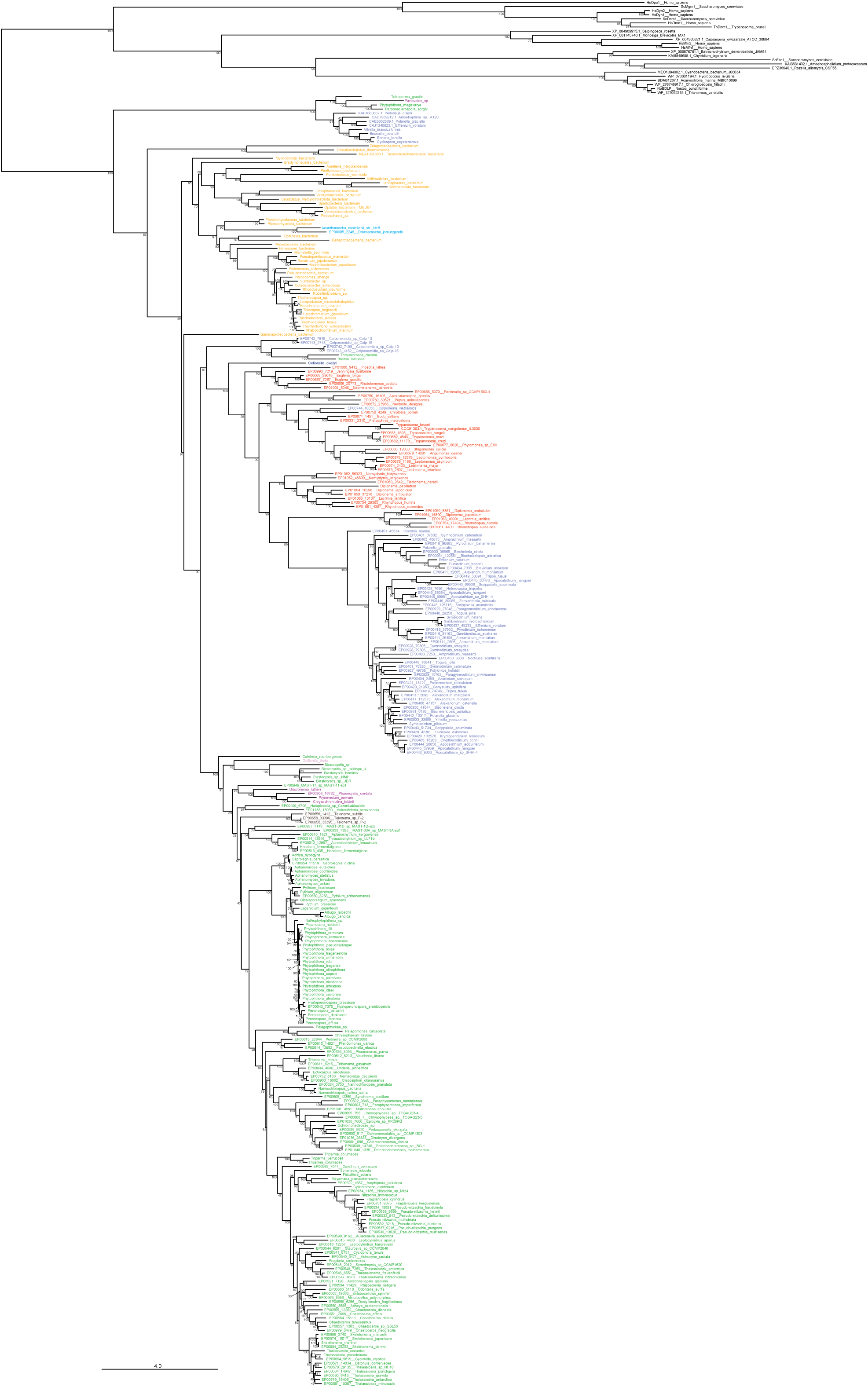
Maximum likelihood phylogenetic tree of eukaryotic and eukaryotic DSPs. The tree was reconstructed using the LG+C60+G4 model of sequence evolution. Ultrafast bootstrap support values (1000 replicates) are shown on each branch. Taxa are colored according to their taxonomic affiliation.

**Figure S4.**
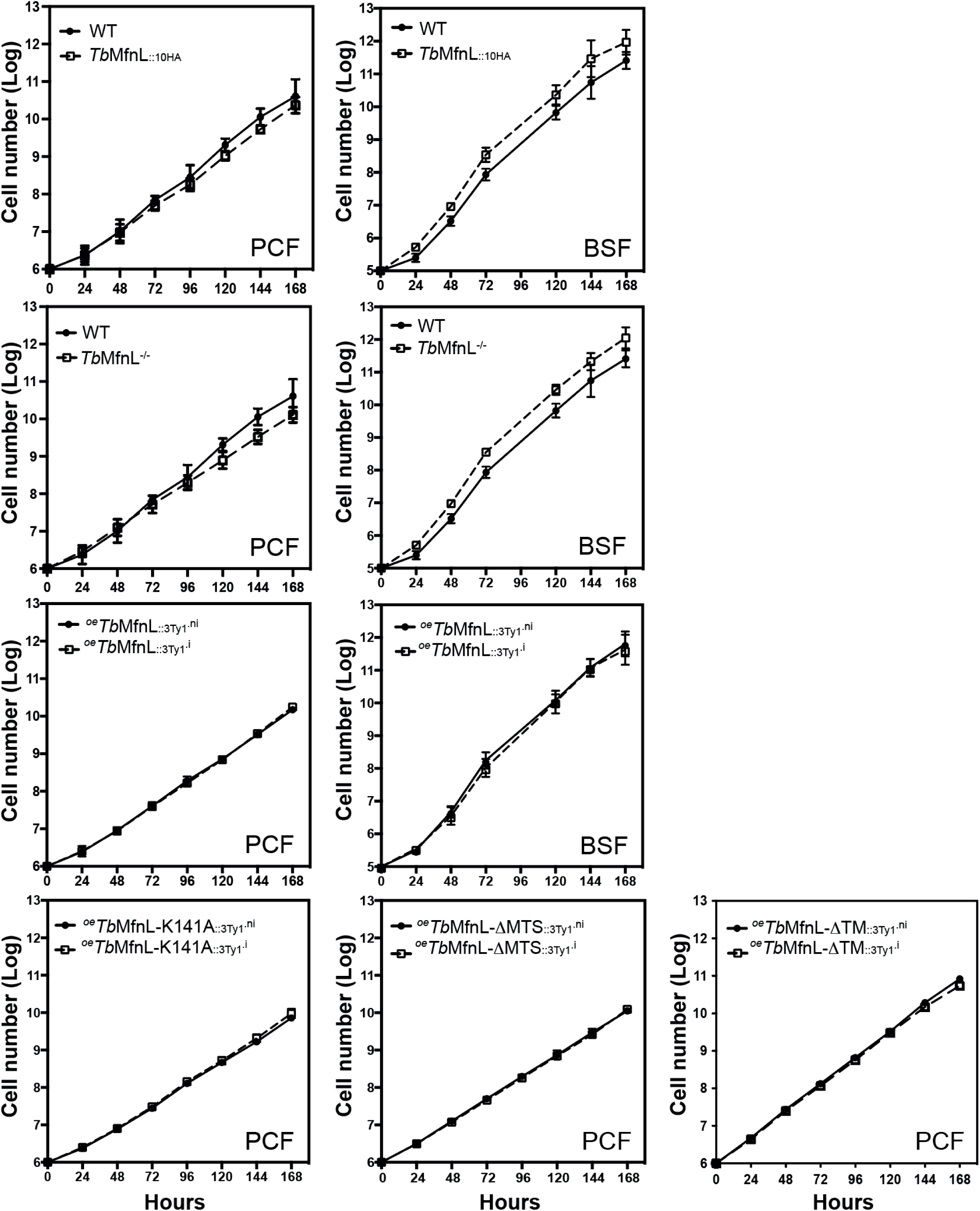
Growth curves of parental, over-expressing and mutant cells. Cells were maintained in the exponential growth phase and cumulative cell numbers reflect normalization for dilution during cultivation. (.ni) non-induced and (.i) induced cells. PCF, procyclic forms; BSF, bloodstream forms. n=3 to 6.

**Figure S5.**
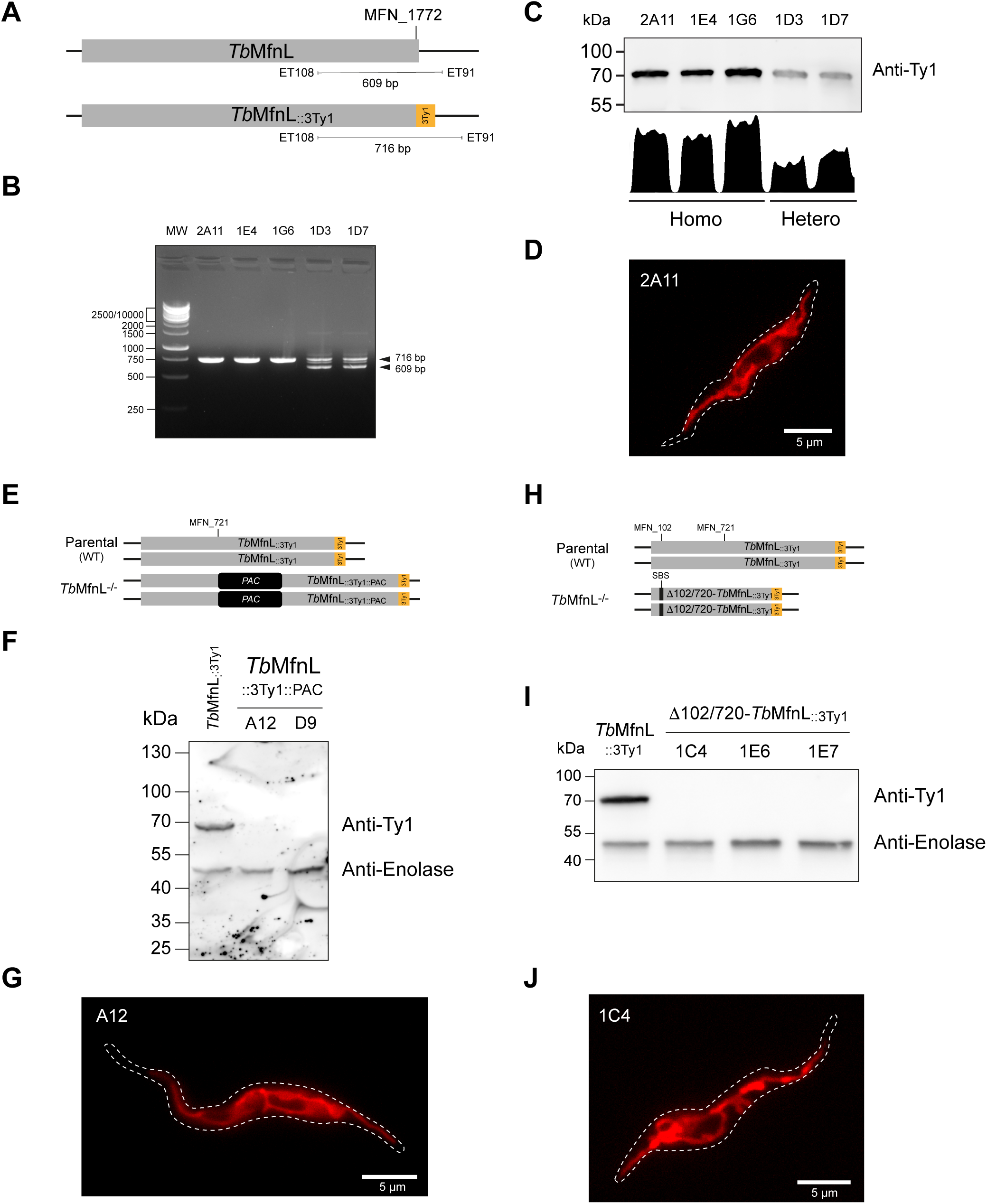
*Tb*MfnL endotagged with a 3xTy1 sequence and inactivation of *Tb*MfnL expression by CRISPR/-Cas9 in PCF. **(A)** Design of the endotagged strain with addition of the 3Ty1 tags at the C-terminus of the protein using the CRISPR/Cas9 system. **(B)** Selection of homozygous clones by PCR (2A11, 1E4 and 1G6) using the PCR approach shown in panel A. **(C)** Western-blot analysis of whole cell extracts (5×10^6^ cells) from *T. brucei* PCF clones expressing *Tb*MfnL_::3Ty1_ revealed with anti-Ty1 antibody and anti-Enolase as a control, and expression levels between homozygous and heterozygous cells. **(D)** Analysis of mitochondrial structure by rhodamine 123 staining in live cells (clone 2A11); **(E)** CRISPR/Cas9-mediated inactivation of *Tb*MfnL using a single guide RNA and insertion of a puromycin resistance marker (PAC) in the *Tb*MfnL_::3Ty1_ (2A11 clone). **(F)** Western blot of whole cell extracts (5×10^6^ cells) of *Tb*MfnL_::3Ty1_ (clone 2A11 background) and *Tb*MfnL_::3Ty1_::PAC (A12 and D9) revealed by anti-Ty1 antibody. **(G)** Rhodamine staining of *Tb*MfnL_::3Ty1_::PAC (clone A12). **(H)** CRISPR/Cas9-mediated inactivation of *Tb*MfnL using two different guide RNAs that deletes approximately 600 bp of the coding sequence in the *Tb*MfnL_::3Ty1_ (2A11 background). **(I)** Western blot of whole cell extracts (5×10^6^ cells) of *Tb*MfnL_::3Ty1_ (2A11 background) and Δ102/720-*Tb*Mfn-L_::3Ty1_ clones (1C4, 1E6 and 1E7) revealed by anti-Ty1 and anti-Enolase antibodies. **(J)** Rhodamine staining of Δ102/720-*Tb*MfnL_::3Ty1_ (clone 1C4).

**Figure S6.**
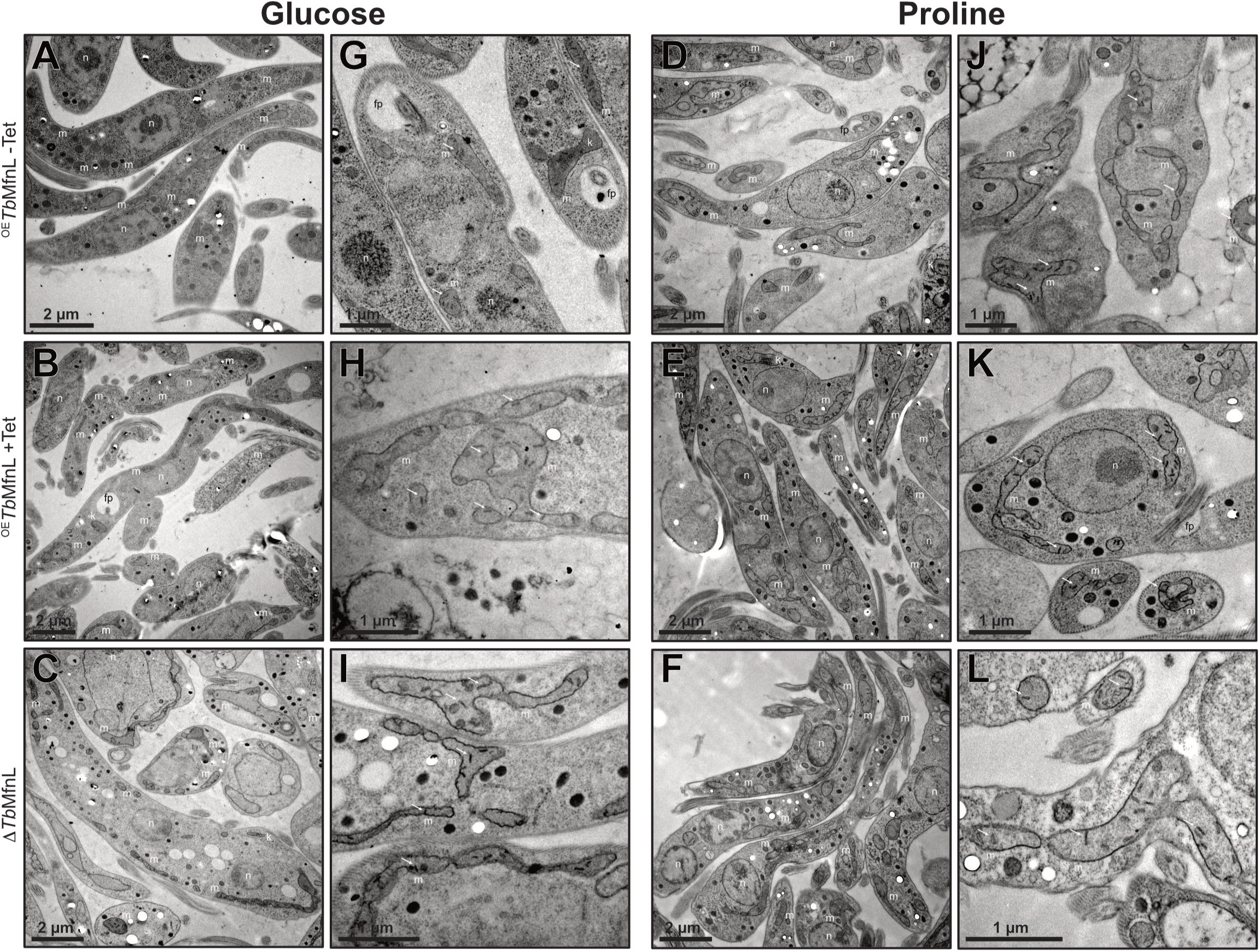
Mitochondrial ultrastructure analyzed by transmission electron microscopy. (TEM). Cells cultured with either glucose or proline, including those overexpressing *Tb*MfnL_::_3Tyl and inactivated TbMfnL^-^/^-^, were examined by TEM. Overexpression of *Tb*MfnL_::_3Tyl was induced for 5 days with tetracycline (Tet) prior to TEM analysis. Panels A-F provide an overview, while panels G-L focus on the mitochondrion and cristaes. n, nucleus; m, mitochondrion; k, kinetoplast; fp, flagellar pocket; white arrows point to the cristae.

**Figure S7.**
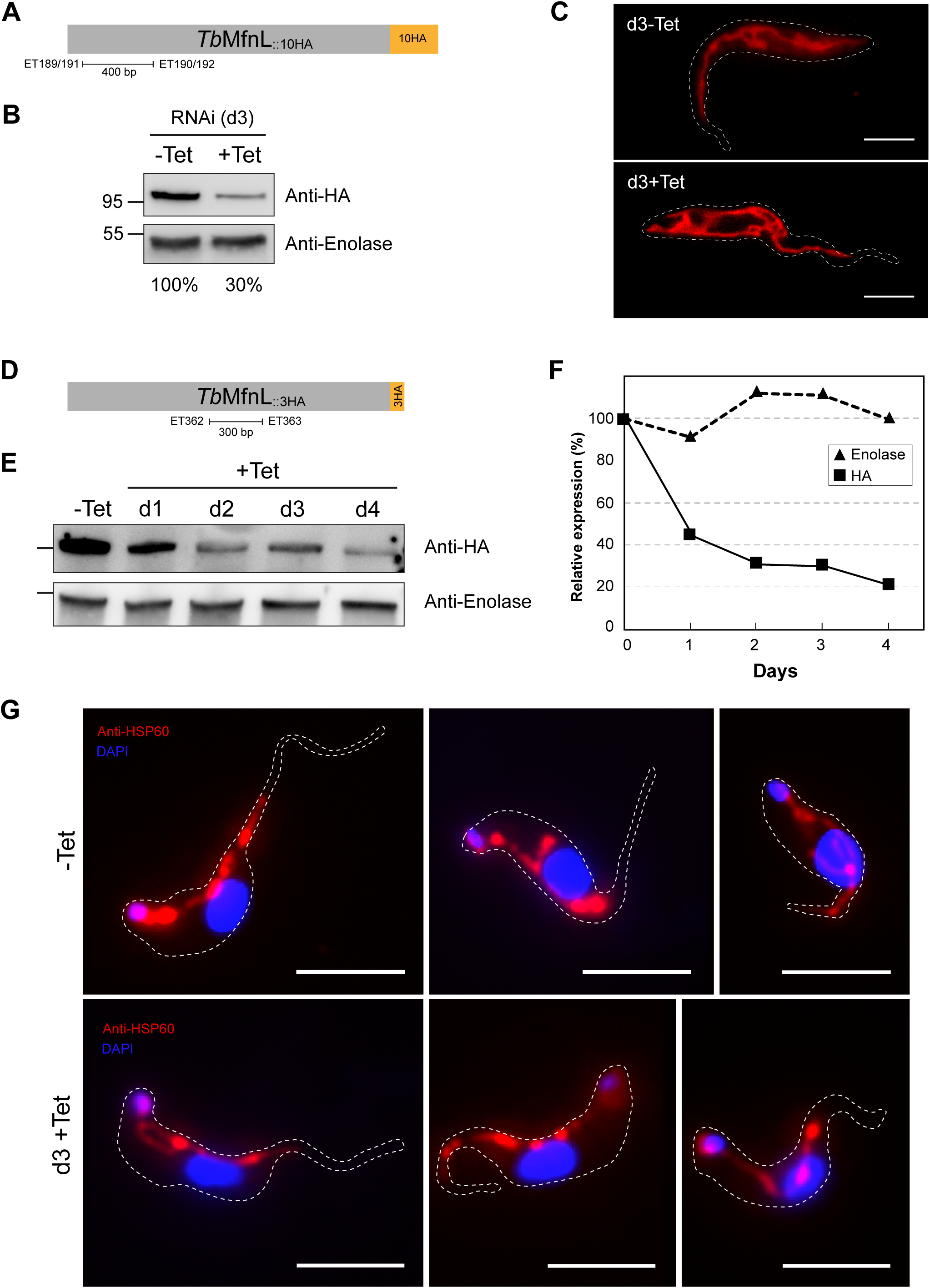
Inhibition of *Tb*MfnL expression by RNAi in PCF (A/B/C) and BSF (D/E/F/G). **(A)** Shematic representation of the sequence used for down-regulation of *Tb*MfnL expression in PCF. **(B)** *Tb*MfnL_::10HA_ down-regulation by RNA interference in PCF. Western blot analysis of total extract (5×10^6^ PCF cells) of non-induced (-Tet) and induced (+Tet) cells using anti-HA antibody after 3 days of tetracycline induction. Antibody against Enolase was used as a loading control. **(C)** Rhodamine 123 labelled mitochondrial network of cells incubated with tetracycline for 3 days. **(D)** Shematic representation of the sequence used for down-regulation of *Tb*MfnL expression in BSF. **(E)** *Tb*MfnL_::3HA_ down-regulation by tetracycline-induced RNA interference in BSF. Western blot analysis of the total extract (5×10^6^ BSF cells) of non-induced (-Tet) and induced (+Tet) cells using anti-HA antibody after 4 days of tetracycline induction. Antibody against Enolase was used as a loading control. **(F)** Relative expression of down-regulation of *Tb*MfnL expression by RNAi in BSF. **(G)** Mitochondrial network labelled with anti-HSP60 from cells incubated or not with tetracycline for 3 days. The scale bar represents 5 μm.

**Figure S8.**
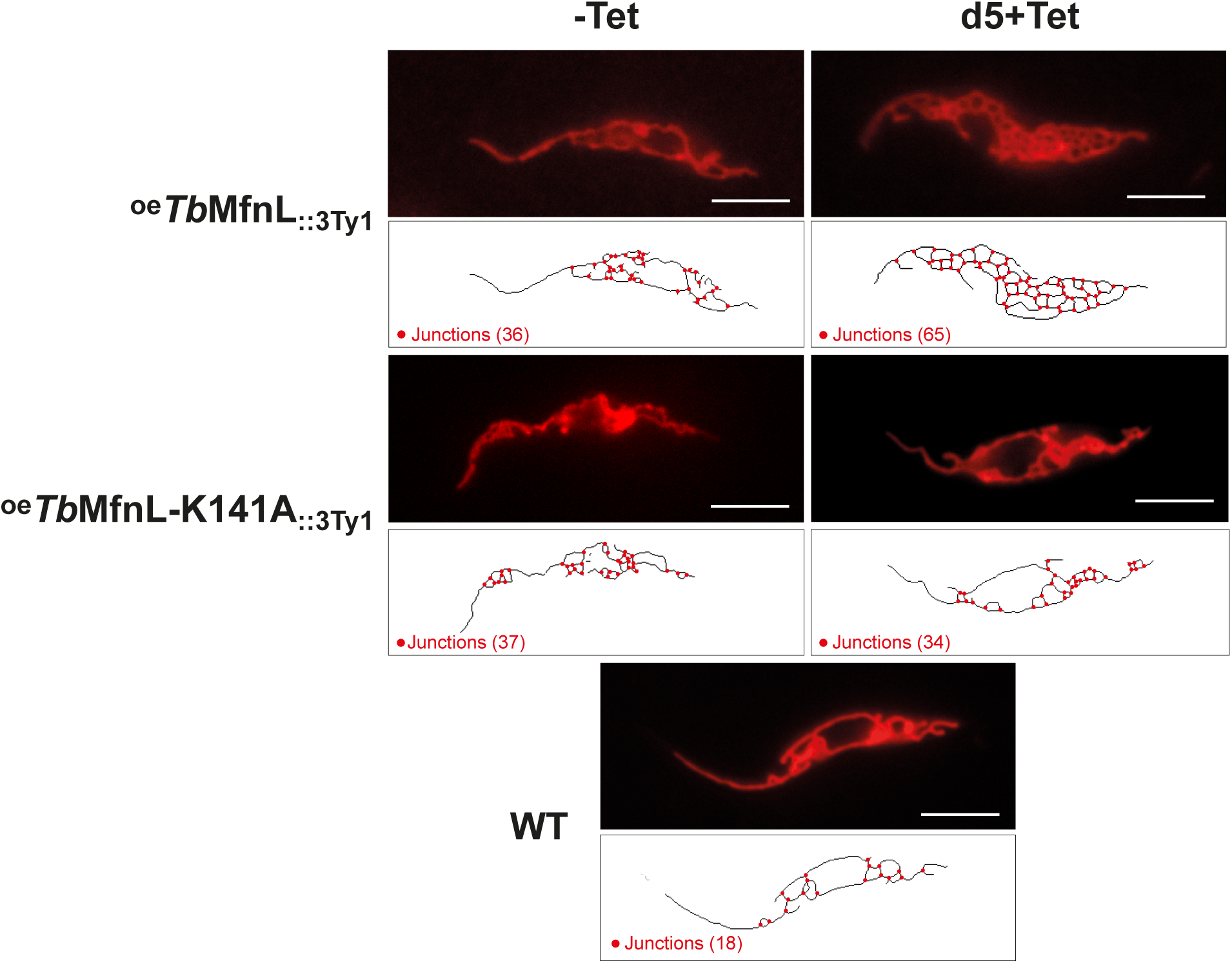
Mitochondrial shape of parental WT, *^oe^Tb*MfnL_::3Ty1_ and *^oe^Tb-* MfnL-K141A_::3Ty1_ PCF cells. The top panel depicts live cells stained with rhodamine 123. Bottom panel was obtained using the ImageJ macro (Mitochondrial Junctions). Tet refers to tetracycline, d5, tetracycline induced cells for 5 days. The scale bar corresponds to 5 µm. ni, non-induced cells; d5 .i, induced cells for 5 days. The scale bar corresponds to 5 µm. Red circles in the skeleton image represent the junctions of three mitochondrial tubules.

**Figure S9.**
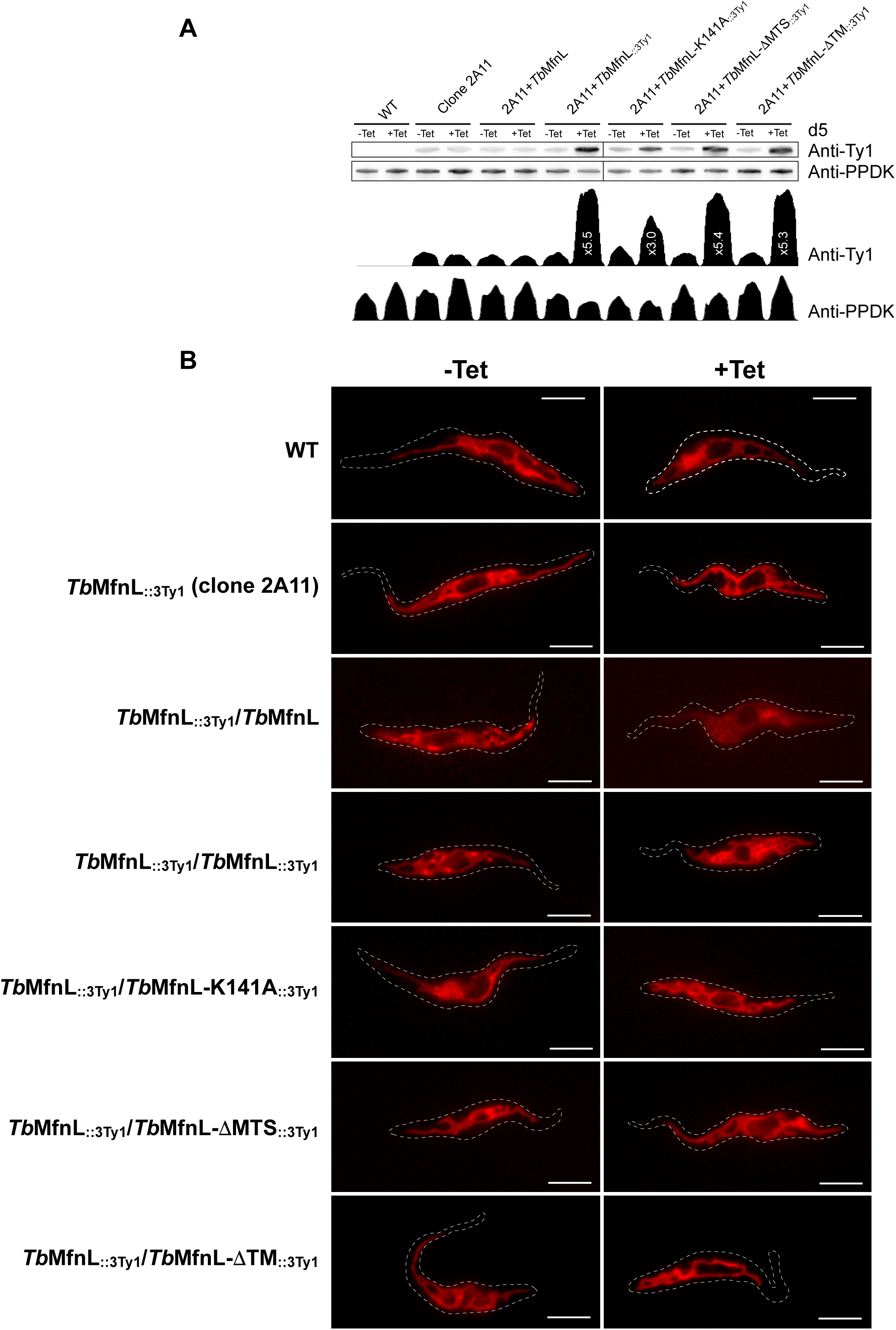
Overexpression of *Tb*MfnL in endotagged cells. **(A)** Western-blot of whole-cell extracts (5×10^6^ cells) of *T. brucei* parental WT PCF and expressing various *Tb*MfnL in *Tb*MfnL_::3Ty1_ background (clone 2A11) revealed with an anti-Ty1 antibody. Quantification of the expression were performed using ImageJ with PPDK as a loading control. **(B)** Mitochondrial structure analysis using rhodamine 123 staining on living cells; mitochondrial structure of parental WT PCF and expressing various *Tb*MfnL in *Tb*MfnL::3Ty1 background without tetracycline (-Tet) and 5 days after addition of tetracycline (+Tet). The scale bar represents 5 μm.

**Figure S11.**
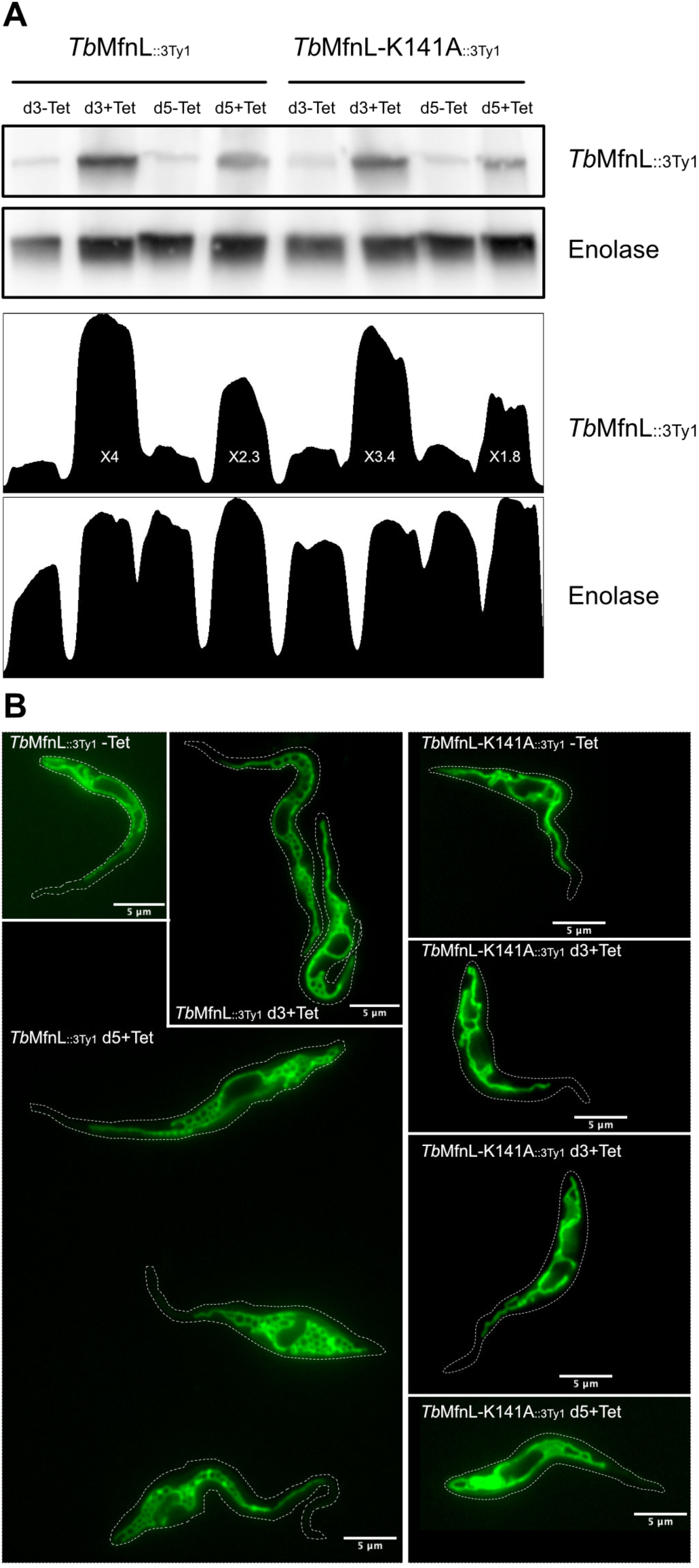
Immuno localization of *^oe^Tb*MfnL_::3Ty1_ by electron microscopy. Immuno localization of *^oe^Tb*MfnL_::3Ty1_ **(A)** and *^oe^Tb*DBFK141A_::3Ty1_ **(B)** expressed in PCF by electron microscopy. Non-induced cells (-Tet, a and b) and 5 days induced cells (+Tet, c and d) were labelled with an anti-Ty1 antibody and an anti-mouse IgG-gold particle conjugate; black dots represent gold particles highlighted by white arrows; b/d are magnification of a and c, respectively. **(C)** Quantification of mitochondrial gold particles. For non-induced (-Tet) and induced (+Tet) *^oe^Tb*MfnL_::3Ty1_ cells, 221 and 3076 gold particles were counted, respectively, and for non-induced (-Tet) and induced (+Tet) *^oe^Tb*MfnL-K141A_::3Ty1_ cells, 78 and 1123 gold particles were counted, respectively. A mean of the non-induced (-Tet) cells from the two strains is represented on the graph. Statistic: t test - Confidence interval 95% - P-value style: 0.1234 (ns); 0.0332 (*); 0.0021 (**); 0.0002 (***); <0.0001 (****).

**Figure S10.**
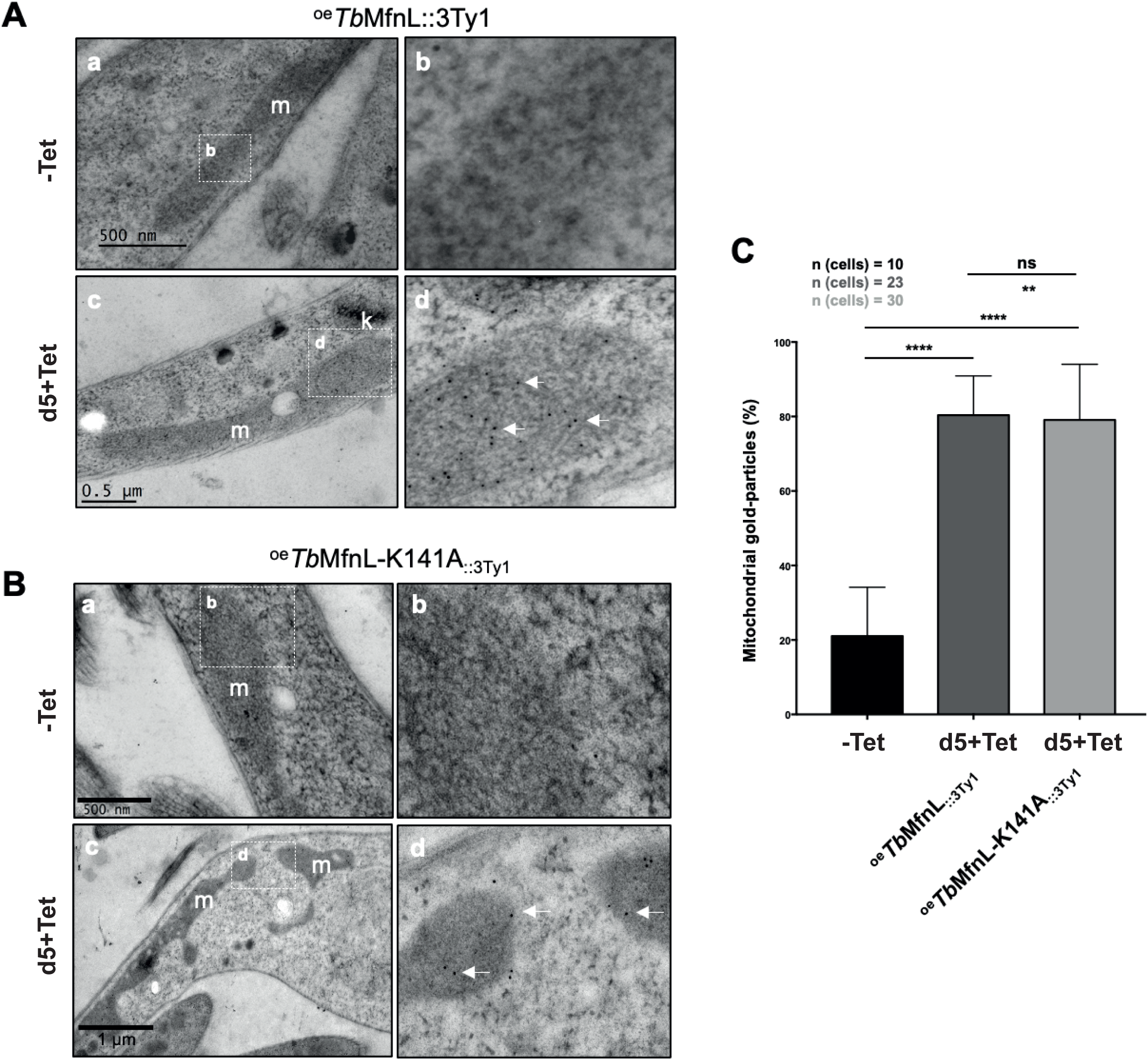
Variation in the expression level of *Tb*MfnL_::3Tyi_ induced by tetracycline.. (A) Western blot analysis of whole-cell extracts (5 × 10^6^ cells) from *T. brucei* PCF expressing *Tb*MfnL_::3Tyi_ and *Tb*MfnL-141A::3Tyi in the *Tb*MfnL_::3Tyi_ background (clone 2A11) probed with an anti-Ty1 antibody. Expression levels were quantified using ImageJ, with PPDK as a loading control, at day **3** and day 5 following tetracycline induction. (B) Corresponding mitochondrial structures were visualized using Mitotracker staining on live cells. The scale bar represents 5 µm.

**Figure S12.**
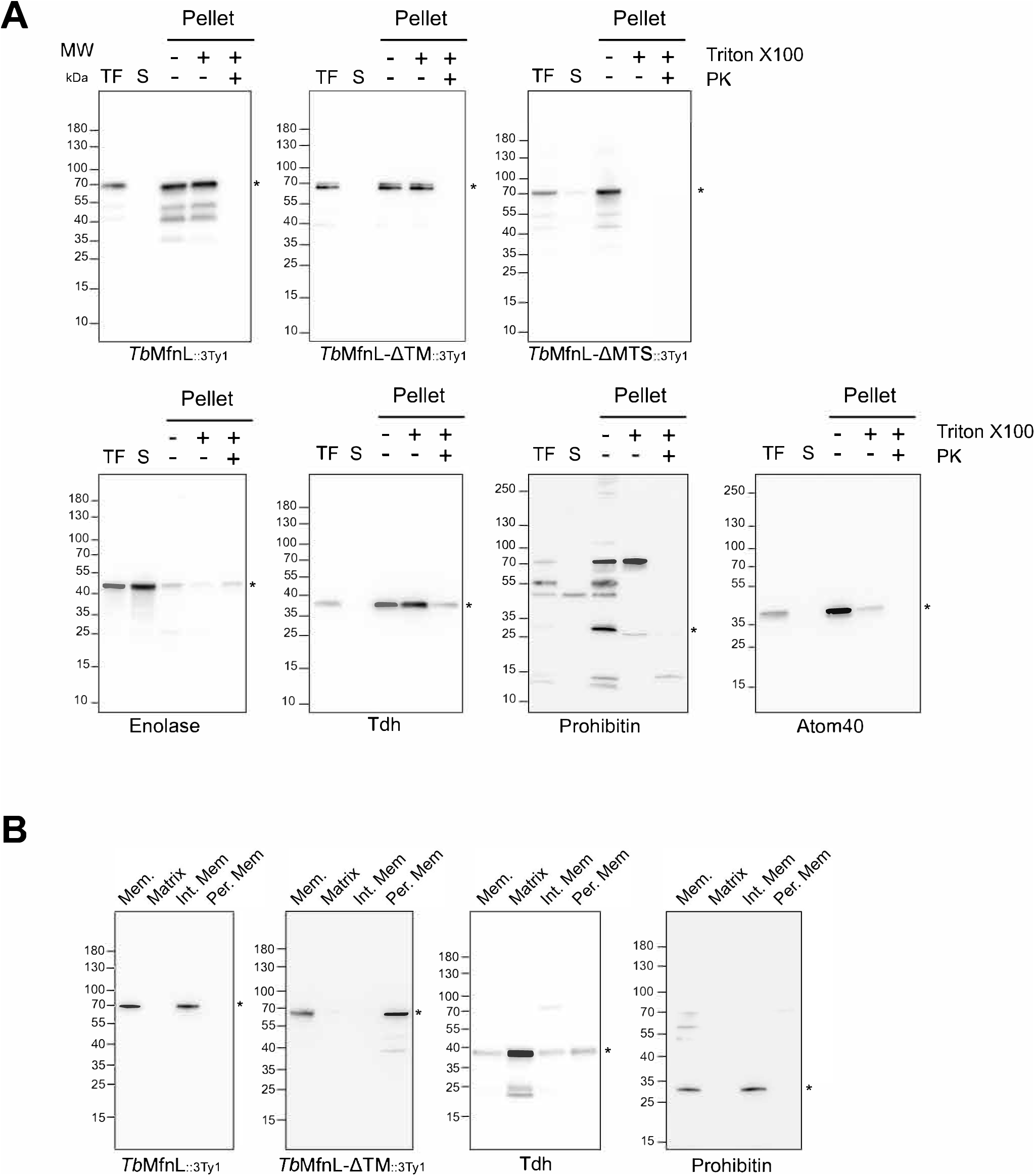
Membrane topology of *Tb*Mfnl (entire western blot). **(A)** The Proteinase K protection assay involved incubating mitoplasts with Proteinase K and Triton X100, as denoted by the“+” symbol. TF represents the total fraction containing both cytoplasmic and membrane components; S denotes the supernatant, comprising the cytosolic contents; and Pellet represents the organelle fraction. **(B)** Carbonate extraction of mitochondria isolated from ^oe^*Tb*MfnL _:: 3Ty1_ and ^oe^*Tb*MfnL-ΔTM_::3Ty1_ cells. Mem., membrane fraction; Matrix, matrix fraction; lnt. Mem, integral membrane fraction; Per. Mem, peripheral membrane fraction. The star indicates the corresponding protein. The expected sizes of the proteins are as follows: *Tb*MfnL - 62 kDa, Enolase - 46 kDa, Tdh - 37 kDa, Prohibitin - 31 kDa, and Atom40 - 38 kDa.

**Figure S13.**
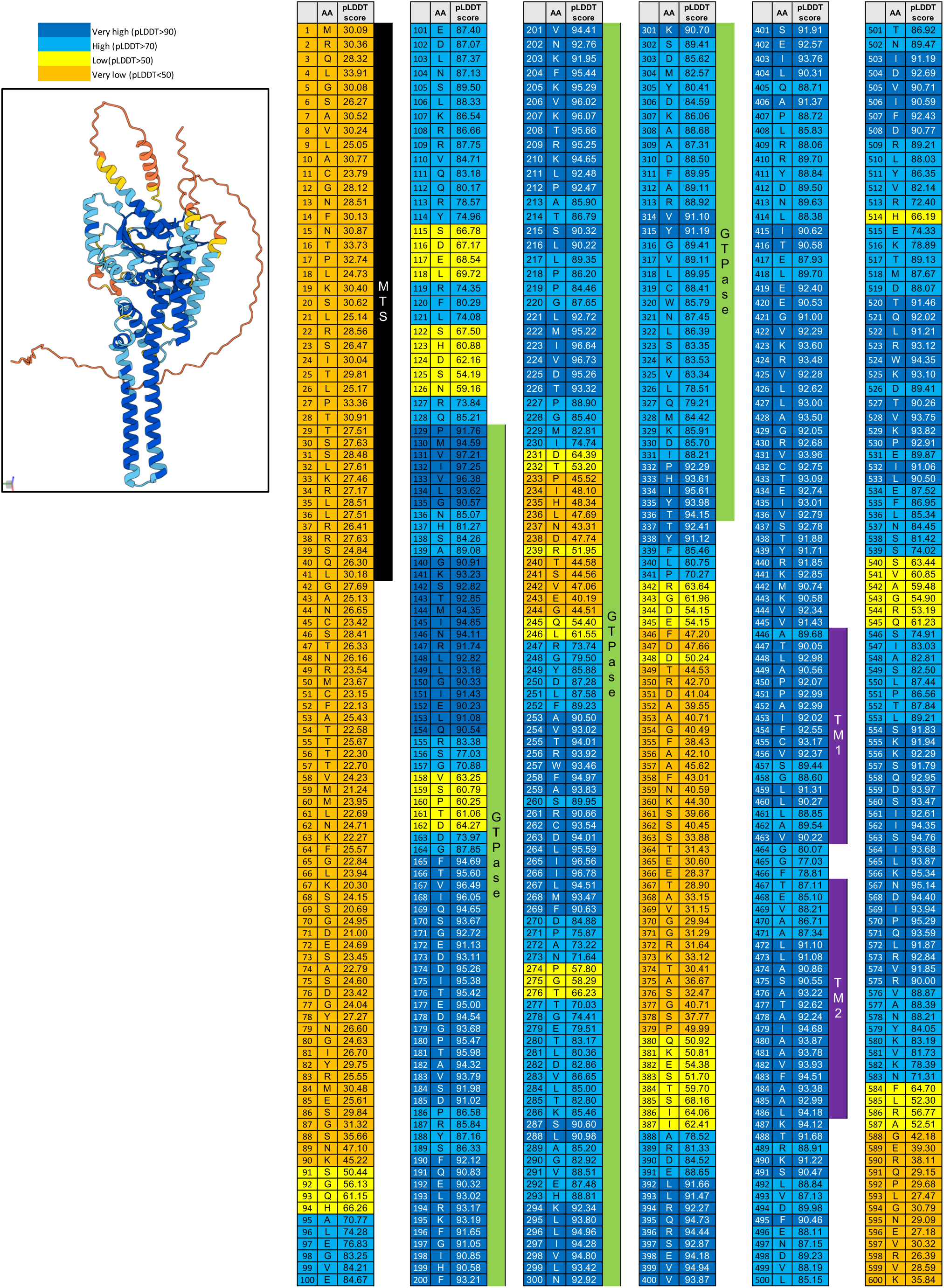
AlphaFold structure of *Tb*MfnL with pLDDT scores for each amino acid. The pLDDT scores are color-coded as blue (>90), light blue (>70), yellow (>50), and orange (<50). The MTS sequence is highlighted by a black box, the GTPase domain by a green box, and the transmembrane domains by purple boxes. The average pLDDT score for the entire protein was 72.5.

